# Gene duplication drove functional divergence of two effectors in the maize anthracnose pathogen

**DOI:** 10.1101/2025.11.26.690860

**Authors:** Pablo García-Rodríguez, Lucía Rodríguez-Mónaco, Francisco Borja Cuevas-Fernández, Sioly Becerra, Silvia Gutiérrez-Sánchez, Andrés Jerez-Vanegas, Flávia Rogério, Jossué Ortíz-Álvarez, Riccardo Baroncelli, Eduardo Antonio Espeso, Michael R. Thon, Serenella A. Sukno

**Affiliations:** Universidad de Salamanca – Instituto de Investigación en Agrobiotecnología (CIALE), Villamayor (Salamanca), Spain; Programa “Investigadoras e Investigadores por México” - Secretaría de Ciencia, Humanidades, Tecnología e Innovación (SECIHTI) Ciudad de México, Mexico; Università di Bologna, Dipartimento di Scienze e Tecnologie Agro-Alimentari, Bologna. Italy; Centro de Investigaciones Biológicas (CIB) Margarita Salas – Consejo Superior de Investigaciones Científicas (CSIC), Departamento de Biología Molecular y Celular, Madrid, Spain; Department of Plant Pathology, University of Florida, Gainesville, Florida 32611, USA

**Keywords:** *Colletotrichum graminicola*, anthracnose, effector, nucleomodulins, effector evolution, gene duplication, neofunctionalization

## Abstract

*Colletotrichum* species rank among the most important fungal pathogens, threatening food security by infecting nearly all major crops worldwide. *Colletotrichum graminicola*, the causal agent of maize anthracnose, secretes effector proteins to manipulate host defences and promote colonization. Building on previous work characterizing the nuclear effector CgEP1, we characterized its paralog CgEP4, which is highly conserved across strains of *C. graminicola*. Phylogenetic analysis of the two genes and their homologs in other species revealed that they originated from a gene duplication event approximately 28 to 18 million years ago, predating the diversification of the *Graminicola* species complex. This timing aligns with the ecological expansion of C4 grasses, suggesting that the functional divergence of these effectors was an adaptive response to facilitate the colonization of emerging monocot hosts. Functional characterization using gene deletion mutants demonstrated that *CgEP4* has a critical role in pathogenicity, characterized by a significant reduction in virulence, delayed penetration, enhanced papilla formation, and decreased fungal biomass. This virulence defect is associated with compromised host colonization and a failure to suppress basal host defences. In the absence of *CgEP4*, the pathogen also showed defects in general fungal physiology and stress tolerance. Overall, our findings establish *CgEP4* as a new, essential nuclear-localized effector that promotes fungal entry and colonization by manipulating host responses. Our findings demonstrate that evolutionary analysis is a valuable tool for discovering new genes important for host adaptation and pathogen evolution.

## Introduction

The filamentous fungal genus *Colletotrichum* ranks among the ten most critical plant pathogenic fungal groups worldwide because of its broad host range, extensive global distribution, and severe agricultural damage from anthracnose disease (Bhunjun et al., 2021; Dean et al., 2012). The genus currently comprises 340 described species, organized into 20 species complexes, and includes major pathogens of cereals, fruits and legumes (Talhinhas & Baroncelli, 2023). Among these, *Colletotrichum graminicola* causes anthracnose disease in maize (*Zea mays),* which led to major epidemics, including the devastating outbreak in the United States during the 1970s (Bergstrom & Nicholson, 1999).

The geographic distribution of this pathogen has recently expanded, likely driven by human activities and environmental changes. Recent reports of its presence in several countries (Cuevas-Fernández et al., 2019; Gutiérrez-Sánchez et al., 2025; Rogério et al., 2023b; Sanz-Martín et al., 2016b) highlight a potential rise in incidence across Europe (Rogério et al., 2023a). The hemibiotrophic pathogen *C. graminicola* undergoes a two-phase infection process.

During its biotrophic phase, the fungus colonizes living maize cells without visible symptoms and delays host cell death; however, unlike true biotrophs, it does not fully suppress several defence responses which remain active from the earliest hours of infection (Vargas et al., 2012). This is followed by a necrotrophic phase, characterized by host tissue necrosis for nutrient acquisition (Perfect et al., 1999; Vargas et al., 2012). Both phases rely heavily on the secretion of effectors, which are small, secreted molecules – including proteins, metabolites, and RNAs – that manipulate host cell biology to promote infection (Hogenhout et al., 2009; O’Connell et al., 2012; Sun et al., 2023). While traditionally associated with immune suppression, recent perspectives emphasize that fungal effectors can also reprogram host development, metabolism, and even microbiome composition to favour colonization (Ai et al., 2025). Among these, proteinaceous effectors are the most widely studied in plant-fungal interactions. These proteins may act in the apoplast, where they inhibit extracellular host defences, or be translocated into host cells, where they modulate immune signalling, transcriptional activity, and hormone pathways (Dodds & Rathjen, 2010; Lo Presti et al., 2015). Effectors may also be localized to the plant nucleus where they regulate host gene expression and control plant cellular responses for maintaining compatible interactions. In addition, the fast evolutionary diversification of effector proteins between fungal species creates challenges for developing sustainable plant resistance strategies (Seong & Krasileva, 2022).

In recent years, advances in comparative genomics, transcriptomics, and proteomics have significantly improved the identification and functional characterization of fungal effectors, offering new insights into their contribution to host colonization (Wilson & McDowell, 2022). In *Colletotrichum graminicola*, 1474 proteins are predicted to be secreted, but only a small subset has been experimentally characterized and confirmed to function as effectors (Becerra et al., 2023). Among these, the fungalysin-family metalloprotease CgFL significantly contributes to virulence by inhibiting maize chitinases during the biotrophic phase (Sanz-Martín et al., 2016a). Additionally, members of the CgCFEM family, which are small cysteine-rich proteins bearing a CFEM domain, suppress defence-related programmed cell death when transiently expressed in *Nicotiana benthamiana*, suggesting a role in evasion of the plant immune system. (Gong et al., 2020). Other factors implicated in pathogenicity include the gene cluster comprising CLU5a and CLU5d, for appressorium penetration peg formation and subsequent host tissue invasion (Eisermann et al., 2019), as well as endoplasmic reticulum-associated proteins such as CgALG3 and CgCNX1, which mediate proper glycosylation and folding of multiple effectors, ensuring their secretion and stability (Mei et al., 2023).

CgEP1 is the first effector in *C. graminicola* demonstrated to target host nuclei and is essential for virulence (Vargas et al., 2016). During the biotrophic phase, it is secreted and translocated into maize nuclei, where it binds chromatin and interacts with host DNA in a non-sequence-specific manner. The deletion of *CgEP1* leads to impaired penetration and strong activation of basal defences, indicating a role in suppressing early immune responses, thus exhibiting characteristics similar to those of bacterial nucleomodulins. CgEP1 served as the first identified nuclear-targeted effector in *C. graminicola* while providing researchers with methods to discover additional effectors that share similar mechanisms of action (Vargas et al., 2016).

The study by Vargas et al. (2016) found that the N-terminus of CgEP1 closely resembles another protein in the *C. graminicola* genome, GLRG_09337 (now known as CGRA01v4_09797) which we named CgEP4 (*Colletotrichum graminicola* Effector Protein 4). A dot plot comparing CgEP4 and CgEP1 revealed that CgEP1’s repeat units exhibit 36% identity to a segment of CgEP4 (Vargas et al., 2016). This finding suggests that CgEP1 and CgEP4 are paralogs, stemming from a gene duplication in an ancestor of the genus *Colletotrichum*. After this duplication, the structure of the *CgEP1* gene underwent remodelling due to mutations and the intragenic duplication in a region that now constitutes CgEP1’s intragenic tandem repeats. However, one outstanding question is whether CgEP1 gained the capacity to modulate the host’s immune system before or after the gene duplication event. This distinction is important for understanding how effector-encoding genes evolve and how often the capacity to modulate the host’s gene expression evolves. To address this question, it is essential to determine whether the paralog CgEP4 also plays a role as an effector. If so, this would imply that the ancestral gene was likely an effector as well, and despite considerable sequence divergence, both genes have retained their ancestral functions.

In this report, we present an updated phylogenetic analysis that reveals the evolution of this gene family has been shaped by a complex history of duplication and loss events. CgEP1 and CgEP4 originated from a gene duplication event that predates the diversification of the Graminicola species complex. *In planta* localization studies show that CgEP4 is translocated to the host nucleus during the early stages of infection and contributes to pathogen virulence. Additionally, gene deletion experiments demonstrate that its absence triggers stronger immune responses and disrupts metabolic balance. This study uncovers a new layer of the molecular strategies employed by *C. graminicola* to manipulate its host, providing deeper insight into the evolution and the role of nuclear effectors in maize anthracnose. Moreover, it shows that evolutionary analysis is a powerful tool for identifying novel traits essential to host adaptation and pathogen evolution.

## Results

### CgEP4 encodes a predicted secreted nuclear effector

The predicted gene structure of *CgEP4* consists of a 79 bp 5′ untranslated region (UTR) and a coding sequence (CDS) of 468 bp interrupted by a single intron initially predicted to be 30 bp in length (Becerra et al., 2023). To confirm this structure experimentally, we performed RT-PCR amplification and sequencing of the region surrounding the predicted intron. Alignment of the resulting cDNA sequence with the predicted CDS revealed a 30 bp discrepancy, indicating that the intron is actually 60 bp (Figure 1A). The *CgEP4* CDS encodes a protein of 145 amino acids. The signal peptide prediction from SignalP 6.0 (Teufel et al., 2022) showed a Sec/SPI-type signal sequence in the first 25 amino acids, with a likelihood score of 0.9998 and a probability of cleavage of 0.739 between positions 25 and 26 (Figure 1B). The signal peptide sequence indicates that CgEP4 would be secreted through the Sec-dependent pathway by Signal Peptidase I cleavage, thus suggesting extracellular function as an effector protein during host-pathogen interactions (Von Heijne, 1990). Furthermore, the NLStradamus r.9 (Nguyen Ba et al., 2009) prediction tool identified a nuclear localization sequence spanning 12 amino acids with 76.6% confidence. The structural properties of CgEP4 appeared unique because no recognized functional domains were detected. Tertiary structure prediction using AlphaFold2 produced a stable 3D model comprising three α-helices (Figure 1C). The molecular dynamics simulations confirmed that the model remained structurally sound when exposed to physiological conditions, thus validating its potential functionality as a protein.

**Figure 1.**
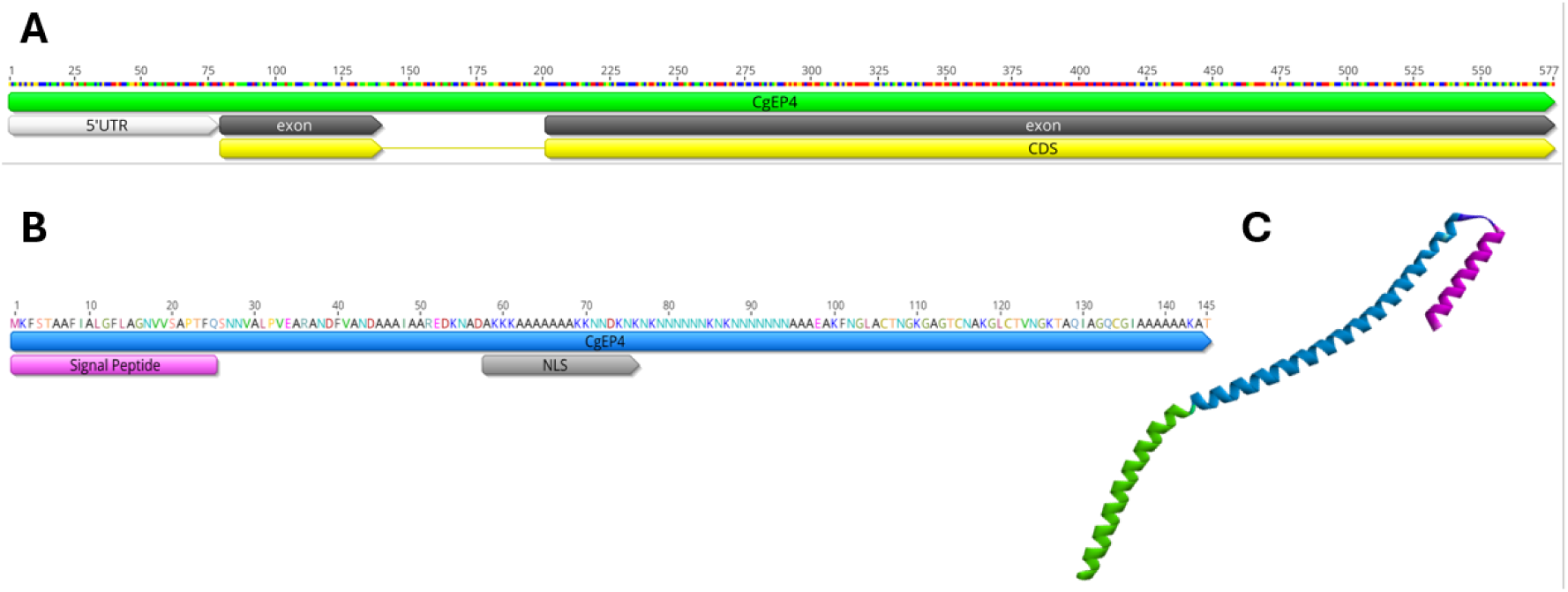
Structural features of CgEP4. **A)** Genomic structure of *CgEP4* (*CGRA01v4_09797*). The 5’ UTR is 79 bp long and the coding sequence (CDS) has two exons (61 and 377 bp) and is divided by an intron of 60 bp. **B)** Primary amino acid sequence of CgEP4. The protein comprises 145 amino acids and includes a predicted N-terminal signal peptide and nuclear localization signal (NLS). **C)** Predicted tertiary structure of CgEP4 by AlphaFold2.

Analysis of the genomes of 61 field isolates from 17 different countries (Rogério et al., 2025) showed 99.6% sequence identity between isolates, indicating a high level of conservation of this gene within *C. graminicola* (data not shown). The predicted secretion signal, along with the nuclear localization sequence, indicates that *CgEP4* encodes a secreted effector protein capable of entering the nucleus during host colonization.

The multiple sequence alignment of CgEP4 and its orthologs from other *Colletotrichum* species revealed that the N- and C-terminal regions are highly conserved, whereas the central domain is rapidly evolving. CgEP4 was detected in 38 of the 41 *Colletotrichum* proteomes analysed. Pairwise identity between *C. graminicola* CgEP4 and its orthologs is ≥ 75 % in the conserved terminal regions but falls below 40 % in the central segment (Figure S1).

### Evolutionary origin and diversification of the CgEP1/CgEP4 gene family

BLASTp searches versus the NCBI nr database revealed that CgEP1 and CgEP4 share no detectable nucleotide similarity outside *Colletotrichum graminicola* and the *Graminicola* species complex, indicating that this pair of effectors represent a lineage-specific innovation within this clade. Moreover, no significant homologs (based on protein sequences) were identified outside the *Colletotrichum* genus, suggesting that the entire gene family is restricted to this lineage.

To explore the evolutionary relationships of this gene family, we performed a phylogenomic analysis using 41 publicly available *Colletotrichum* proteomes representing the diversity of the genus, with *Verticillium dahliae* as an outgroup to root the phylogeny. An iterative BLASTp search (E-value ≤ 1e−3) using CgEP1 as a query identified 68 homologous sequences (Table S1), which were aligned and used to reconstruct a maximum-likelihood phylogeny. Across *Colletotrichum* species, the presence and copy number of *CgEP1/CgEP4*-like genes vary considerably. Most species outside the *Graminicola* complex possess zero to two homologs, whereas species within the *Graminicola* complex contain two to four copies, consistent with additional duplication events and reduced gene loss in this lineage. Notably, species of the early-diverging *Orbiculare* complex harbour a single copy located either in one or the other main clade, suggesting reciprocal gene loss following an ancestral duplication event. The resulting tree resolved two well-supported major clades (Figure 2): Clade 1 - containing CgEP1, CgEP4, and their orthologs, is dominated by sequences from the *Graminicola* complex, indicating a lineage-specific expansion within this group. Clade 2 - includes a *C. graminicola* protein clustering with representatives from nearly all *Colletotrichum* species complexes, consistent with an ancient duplication that predates the diversification of the genus. Within Clade 1, two distinct subgroups of *Graminicola* complex proteins were identified, consistent with a second duplication event giving rise to CgEP1 and CgEP4. A more recent duplication was detected in *C. eremochloae*, which carries two paralogous copies of the gene.

**Figure 2.**
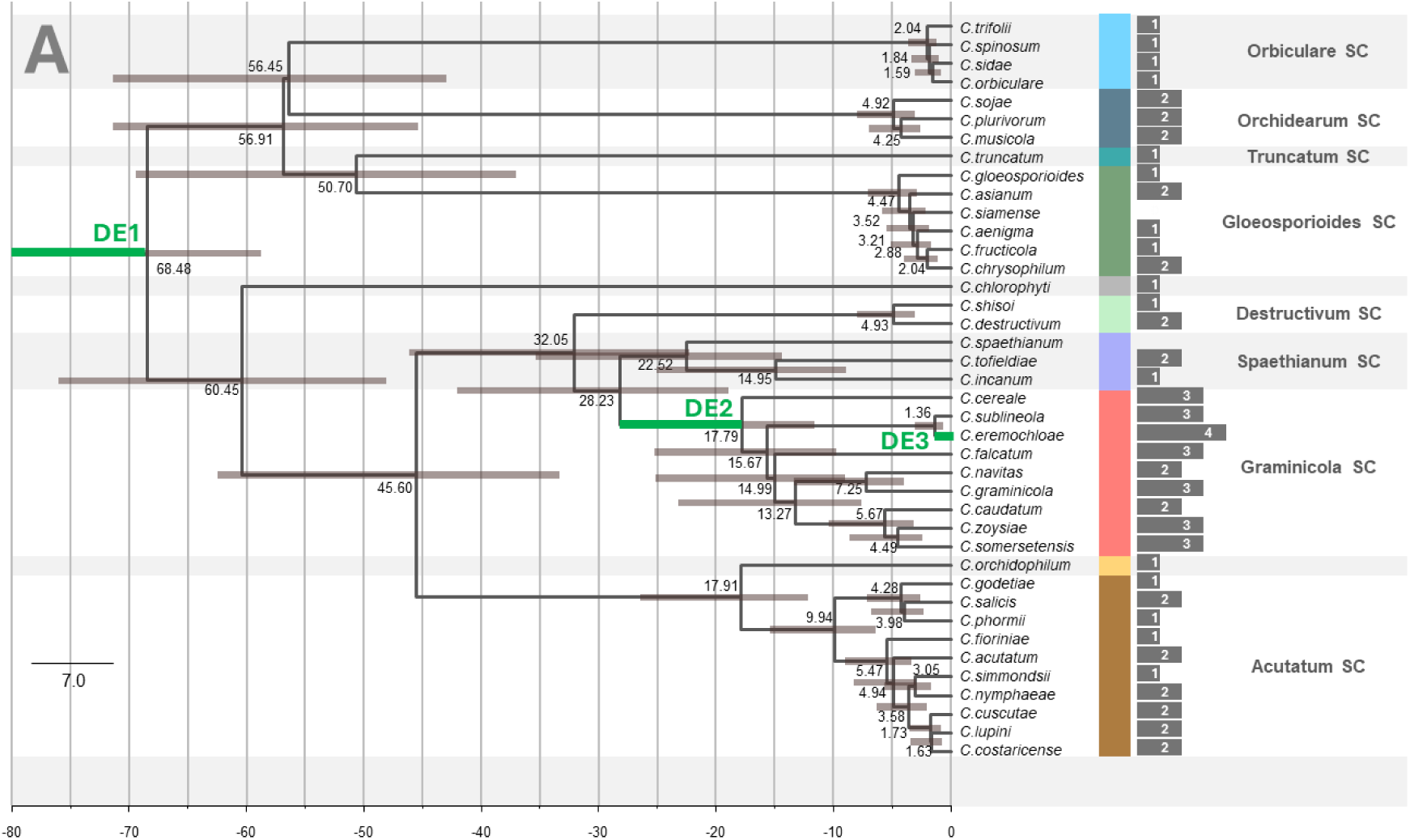
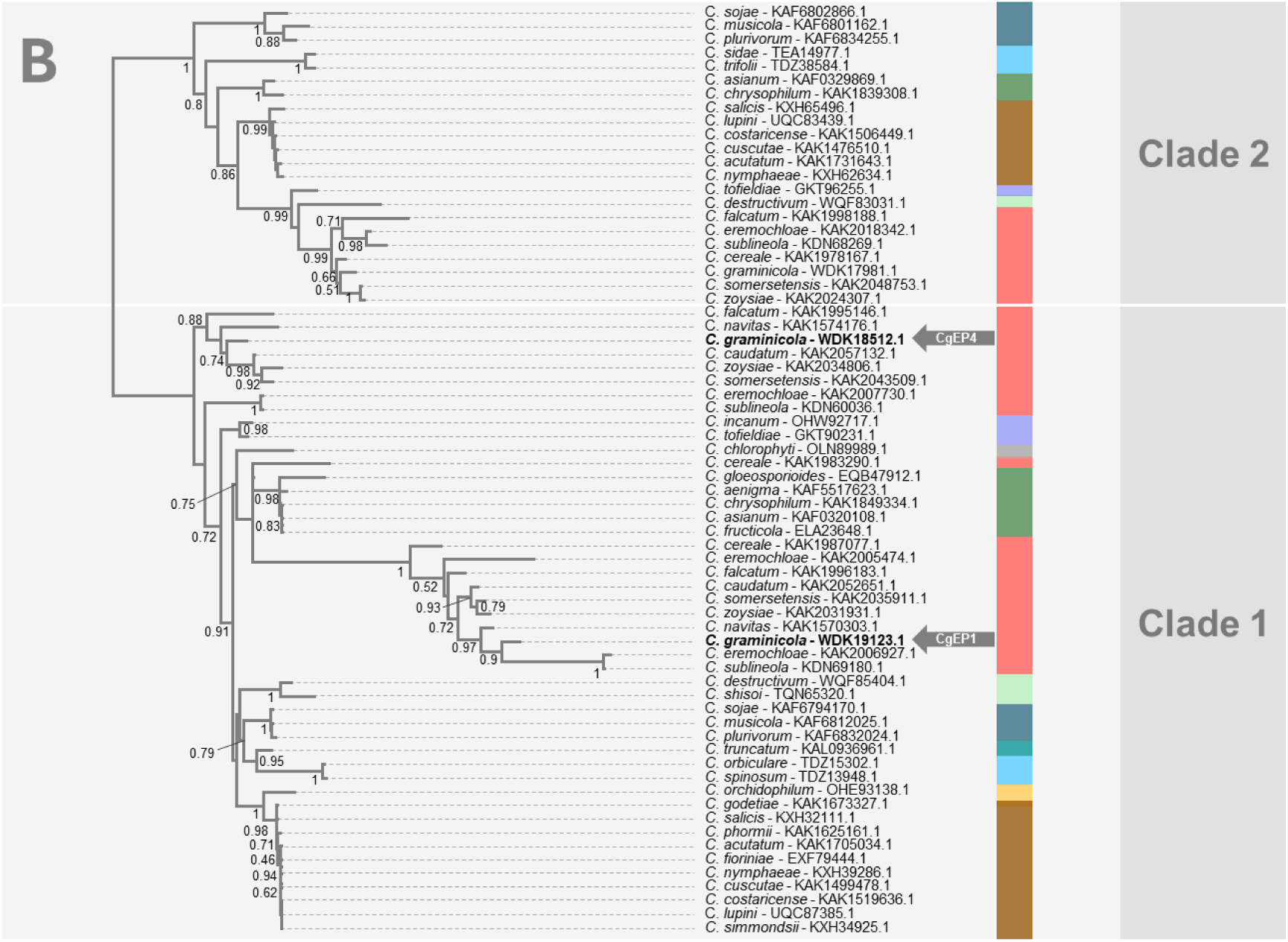
Phylogenetic analysis of the genus *Colletotrichum* and the CgEP1/CgEP4 gene family. **A)** Species-level calibrated phylogeny of 41 *Colletotrichum* species inferred from concatenated alignments of single-copy orthologs identified with OrthoFinder. Tree topology was reconstructed using FastTree and the numbers next to the nodes indicate estimated calibrations expressed in millions of years ago (MYA). Green branches highlight duplication events. Colored bars on the right denote different *Colletotrichum* species complexes; the legend in the lower right indicates the corresponding species complexes. Bar diagrams show the number of CgEP1/CgEP4 and their homologs. *Verticillium dahliae* was used as an outgroup to root the tree. **B)** Maximum-likelihood phylogeny of CgEP1, CgEP4 (indicated by grey arrows), and their homologs identified across 40 *Colletotrichum* proteomes. Numbers next to the node represent support values, while colored bars on the right denote different *Colletotrichum* species complexes.

Calibrated phylogenomic analyses identified three major duplication events in the evolutionary history of this family: DE1, an ancient duplication occurring before ∼68.5 million years ago (Late Cretaceous), predating the diversification of *Colletotrichum*; DE2, a duplication between 28.2 and 17.8 million years ago (Oligocene–Miocene), corresponding to the origin of CgEP1 and CgEP4; DE3, a recent duplication after 1.36 million years ago (Pleistocene), identified in *C. eremochloae*. The Oligocene–Miocene duplication (DE2) occurred during a period of major ecological transformation marked by the early expansion of C4 grasses, the rise of open grasslands, and the diversification of grass-associated fungi. The clear separation of CgEP1 and CgEP4 into distinct, well-supported clades supports post-duplication functional divergence, consistent with their distinct yet complementary roles in host colonization and fungal physiology.

In the reference genome assembly of *C. graminicola* (Becerra et al., 2023), *CgEP4* and its paralog *CgEP1* are both located on chromosome 6 (Figure S2). CgEP4 is encoded within a genomic segment enriched in repetitive elements. The closest short repetitive elements are located 4207 bp upstream and 5612 bp downstream of the CDS, whereas the nearest long terminal repeat (LTR) retrotransposons lie 23 kb upstream and 91 kb downstream. Similarly, CgEP1, encoded on the forward strand in another chromosomal region, is also flanked closely by short repeats at distances of 3153 bp upstream and 5596 bp downstream and the nearest LTR retrotransposon is 52 kb upstream. The proximity of both effector genes to these repeat-rich regions and LTR retrotransposons strongly suggests that transposon-mediated mechanisms and recombination events may have influenced their duplication, genomic rearrangement, and subsequent functional divergence.

### CgEP4 is translocated to maize nuclei during early infection

To predict the subcellular localization of CgEP4 during maize infection, we ran several *in silico* tools. Nuclear localization prediction programs suggested nuclear or nucleolar targeting, while EffectorP 3.0 (Sperschneider & Dodds, 2022) classified it as a likely apoplastic or cytoplasmic effector lacking transmembrane domains (Table S2). To experimentally test these predictions, we generated a C-terminal translational fusion of CgEP4 with mCherry under the control of its native promoter (*CgEP4*-*mCherry*) and transformed it into *C. graminicola* strain M1.001 (Figure S3). PCR confirmed successful integration, and the fluorescent signal was analysed by fluorescence microscopy. At 36 hours post-inoculation (hpi), infected maize leaf sheaths exhibited red fluorescence within host nuclei (Figure 3). Nuclear localization was confirmed by colocalization with DAPI staining. Notably, fluorescence intensity was frequently stronger in a subnuclear compartment consistent with the nucleolus.

**Figure 3.**
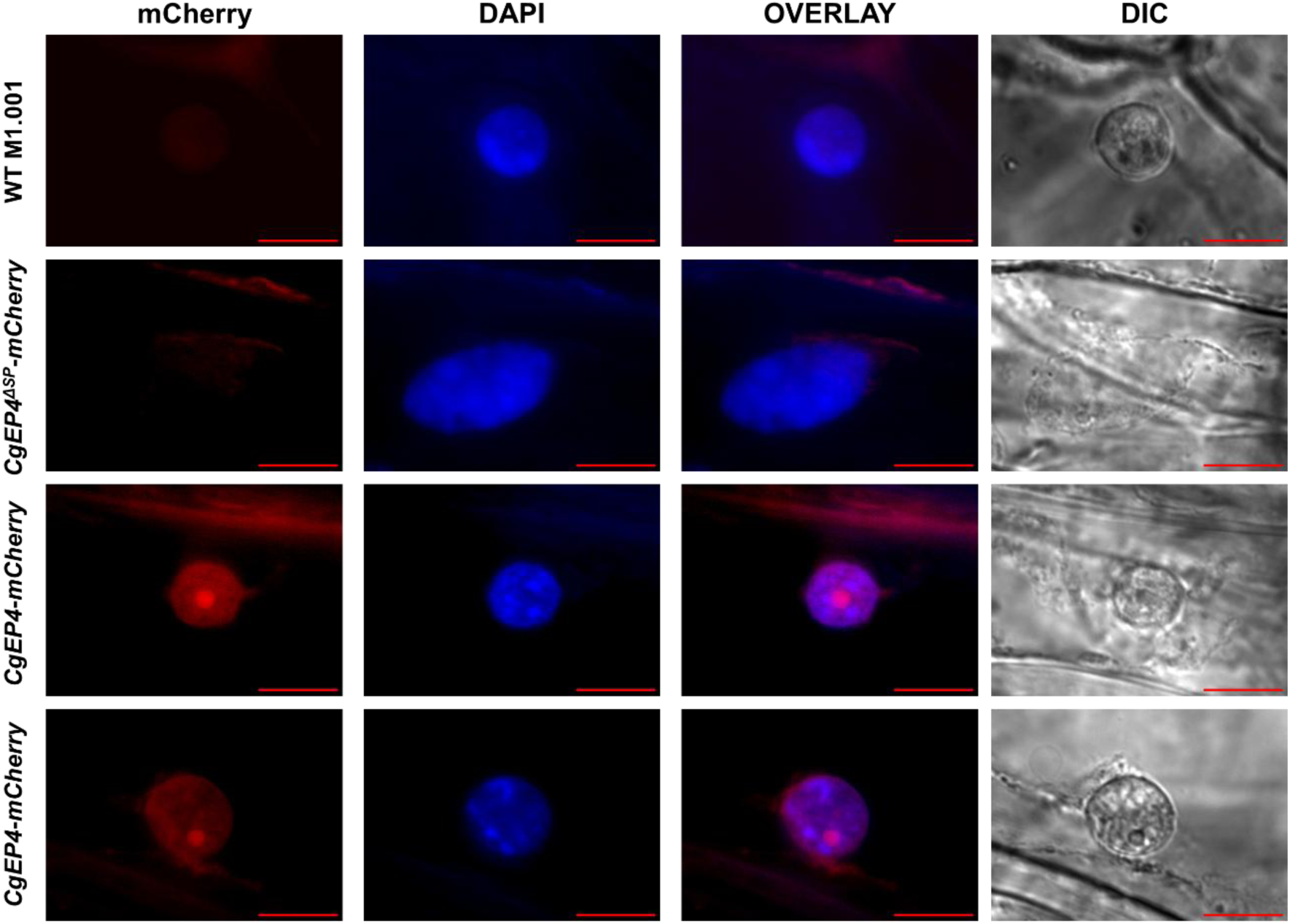
Subcellular localization of CgEP4 during maize infection. CgEP4 fused to red fluorescent protein (mCherry) shows clear nuclear localization in infected maize cells, as confirmed by colocalization with the nuclear stain DAPI. Within the nucleus, mCherry fluorescence was consistently enriched in a defined subnuclear region consistent with nucleolar localization. No nuclear fluorescence was observed in maize cells infected with the wild-type M1.001 strain or with the CgEP4^ΔSP^-mCherry control lacking the signal peptide. Images were acquired at 36 hours post-inoculation (hpi). Scale bar = 20 μm.

To assess the role of secretion, we constructed a truncated version of CgEP4 lacking its signal peptide (*CgEP4*^Δ*SP*^-*mCherry*). In the maize leaf sheath infections with this strain, mCherry fluorescence was restricted to fungal structures such as hyphae conidia and appressoria without showing any host cell signal (Figure 3). The results indicate that secretion is essential for CgEP4 to move into host cell nuclei. Collectively, the results demonstrate that CgEP4 is secreted by the fungus and targets the host cell nucleus during the initial phase of maize infection.

We also expressed a constitutive GFP fusion lacking the signal peptide (*PgpdA::CgEP4^ΔSP^–eGFP*) to examine protein distribution in fungal hyphae grown *in vitro*. Fluorescence was clearly visible within the cytoplasm of vegetative hyphae but absent from the surrounding medium (Figure S4). These observations indicate that removal of the signal peptide prevents secretion and confirms that the N-terminal signal peptide is functional and necessary for secretion.

### CgEP4 is expressed during the biotrophic phase of infection

*CgEP4* exhibits dynamic transcriptional regulation during maize colonization by *C. graminicola*. The RT-qPCR analysis was performed on mycelium and conidia samples from *in vitro* cultures together with infected maize leaf tissue collected at 24, 36, 48, 60 and 72 hpi and at 4 and 8 days post-inoculation (dpi) following the protocol described in Sanz-Martín et al. (2016a). The examined time points cover the entire biotrophy to necrotrophy transition allowing us to link expression patterns with fungal development *in planta*.

The *CgEP4* gene showed a minimal expression in mycelium and conidia which indicates that its expression becomes mostly active during the infection process. Transcription levels began rising at 24 hours post-inoculation and reached a peak at 48 hours post-inoculation, during the late biotrophic phase, before gradually declining. Gene expression remained detectable throughout the necrotrophic phase at both 60 and 72 hpi and extended into the 4 and 8 dpi time points (Figure 4).

**Figure 4.**
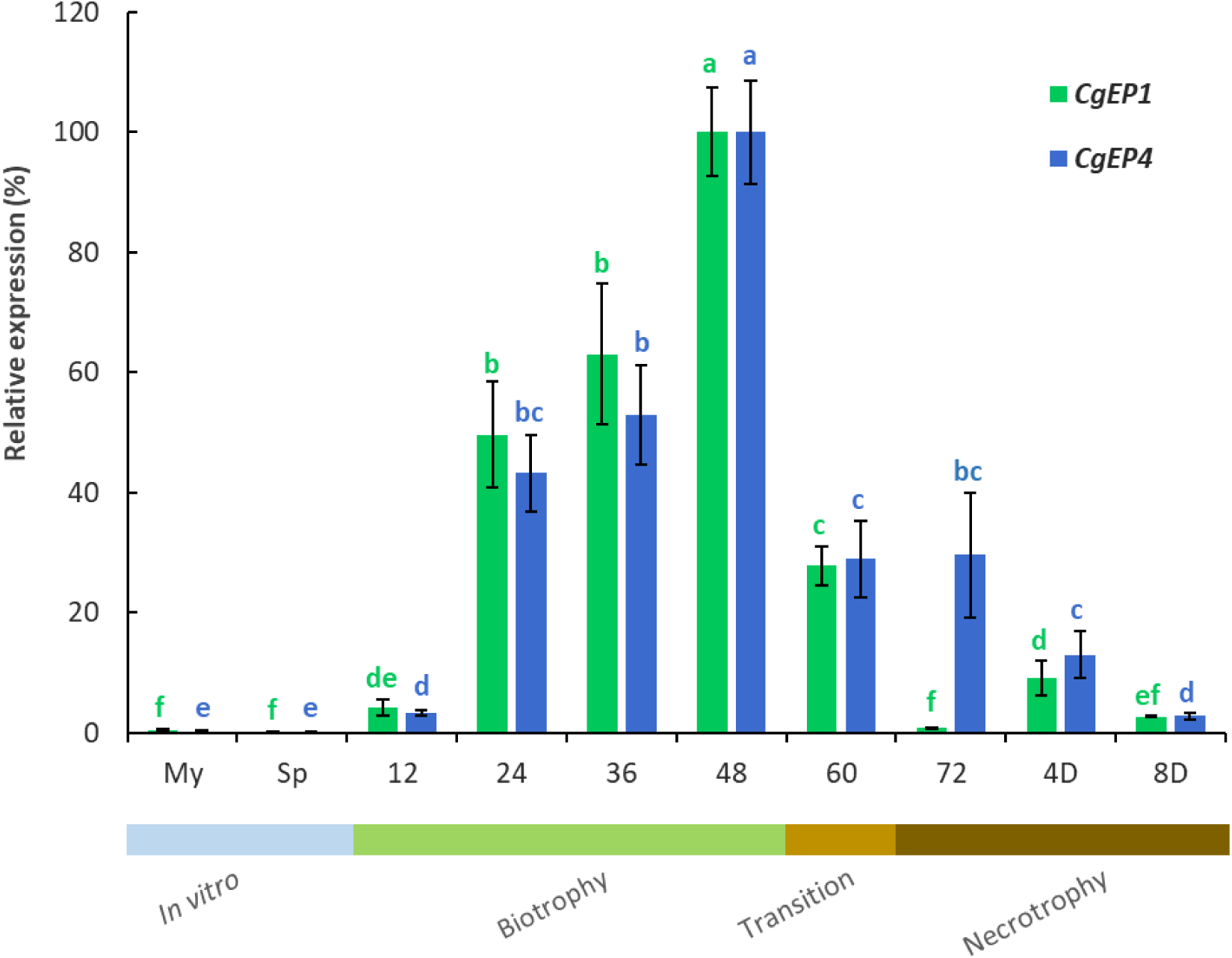
Expression profile of CgEP4. Relative expression of *CgEP4* and *CgEP1* was analysed by RT-qPCR at multiple time points during *C. graminicola* infection of maize leaves, as well as under *in vitro* conditions (My: mycelium; Sp: conidia). Expression levels were normalized to the histone H3 gene and are presented relative to the highest value observed for each gene (set as 100%). Statistical analyses were performed separately for each gene. Bars represent the mean of four biological replicates, each with two technical replicates. Error bars indicate standard deviation. Statistical significance was assessed using the non-parametric Mann–Whitney U test; bars labelled with different letters differ significantly within the same gene (*p* < 0.05). The coloured bar below the graph indicates the approximate infection stage for each time point: *in vitro* (blue), biotrophy (green), transition (light brown), and necrotrophy (dark brown).

To contextualize this expression profile, we compared *CgEP4* with *CgEP1*, a previously characterized effector associated with biotrophic development. Both genes exhibited similar dynamics, with maximal expression at 48 hpi. Notably, *CgEP4* maintained higher transcript levels than *CgEP1* during later infection stages, suggesting a broader temporal window of activity.

The data indicate that *CgEP4* is specifically induced during infection and exhibits its highest expression levels during biotrophy, followed by sustained transcription throughout necrotrophy, supporting its role in establishing host colonization.

### CgEP4 is required for efficient infection and suppression of early host defences

The functional role of *CgEP4* in virulence was investigated by generating two independent knockout strains Δ*CgEP4*-4 and Δ*CgEP4*-6 in *C. graminicola* and testing their virulence on maize aerial tissues. The two mutants produced lesions that averaged 55% smaller than those produced by the wild type M1.001 at 3 days post-inoculation (dpi) in leaf blight assays. We used Δ*CgEP4-6* for further experiments. The phenotype was shown to be caused by gene function loss when *CgEP4* was reintroduced under its native promoter (Δ*CgEP4*::*CgEP4*) (Figure 5A). A comparable reduction in virulence was observed in stalk infection assays, where Δ*CgEP4* strains generated lesions approximately 60% smaller than those of the wild type. The virulence defect was fully rescued by complementation, thus confirming that CgEP4 is necessary for full pathogenicity in both tissue types (Figure 5B).

**Figure 5.**
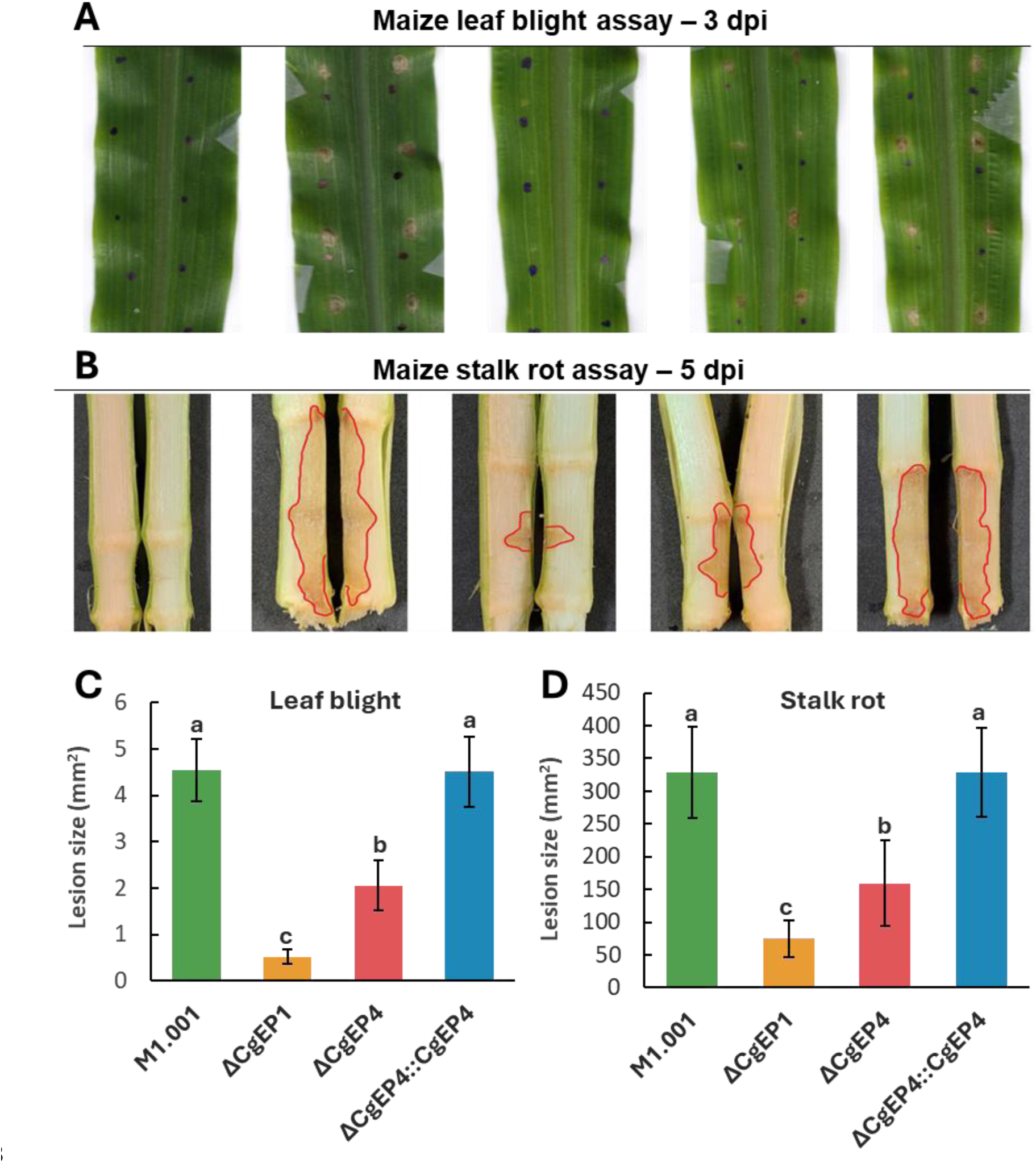
Virulence assays of *C. graminicola* strains on maize leaves and stems. **(A)** Representative leaf blight symptoms at 3 days post-inoculation (dpi). Black dots mark inoculation points. **(B)** Representative stalk rot symptoms in longitudinally split stems at 5 dpi. **(C)** Leaf blight lesion area was quantified for wild-type, Δ*CgEP4*, and strains. Bars represent mean lesion area ± standard deviation (SD). The assay included four biological replicates, each with four technical replicates; 20 infection sites were evaluated per replicate (n = 320 per strain). Statistical analysis was performed using one-way ANOVA followed by Tukey’s HSD test. **(D)** Stalk rot necrotic lesion size was measured in longitudinally split stems following inoculation at the first internode. Bar plots show mean lesion area ± SD (n = 20 per strain). Statistical analysis was performed using one-way ANOVA followed by Tukey’s HSD test (*p < 0.05*). Representative images of symptomatic tissue are shown above each graph.

The qPCR analysis of fungal biomass in infected leaves at 3 dpi showed that Δ*CgEP4* mutants accumulated significantly less fungal DNA than the wild-type strain (Figure S5), indicating that the reduced lesion size may reflect compromised host colonization.

Analysis of the infection process revealed that CgEP4 contributes to efficient host penetration. At 36 hpi, only 5% of Δ*CgEP4* appressoria successfully penetrated the epidermis, compared to ∼70% for M1.001. This defect was comparable to that observed in Δ*CgEP1* mutants (7% penetration), suggesting that both effectors play roles in penetration of the epidermis a particularly critical stage early in infection. The penetration efficiency of Δ*CgEP4* mutants reached 50% at 48 hpi, thereby reducing the difference with the wild type (Figure 6A).

**Figure 6.**
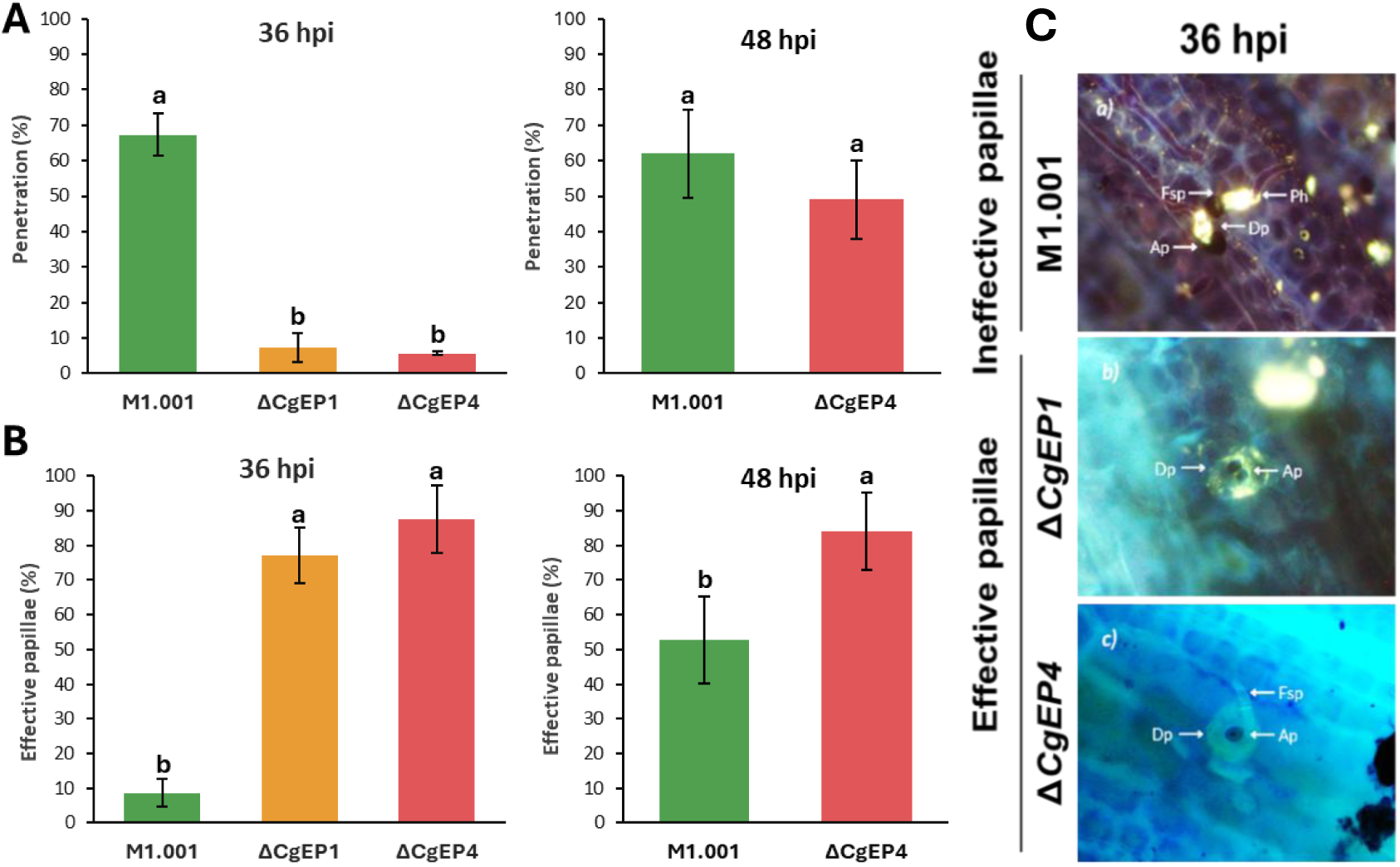
CgEP4 is required for efficient colonization and suppression of host defences. **(A)** Penetration efficiency of appressoria at 36- and 48-hours post-inoculation (hpi), expressed as the percentage of appressoria that successfully penetrated host epidermal cells. **(B)** Frequency of effective papillae at 36 and 48 hpi, defined as papillae that successfully blocked fungal entry (n = 100 appressoria per replicate). Bars represent mean values ± standard deviation from three independent biological replicates. Statistical significance was assessed using a two-tailed Student’s *t*-test; letters indicate significant differences (*p* < 0.05). **(C)** Representative micrographs showing ineffective and effective papillae at appressorial penetration sites. *Dp*: papilla deposition; *Ap*: appressorium; *Fsp*: conidium; *Ph*: primary hypha.

We further examined the host’s localized defence response by quantifying the effectiveness of papilla formation at appressorial contact sites. At 36 hpi, 87% of papillae in Δ*CgEP4*-infected tissues effectively blocked entry, compared to 78% for Δ*CgEP1* and only 8.5% for the wild type. At 48 hpi, papilla resistance remained elevated in Δ*CgEP4* infections (84%) compared to M1.001 (52%), indicating that CgEP4 is necessary to suppress this basal defence response during the early and intermediate stages of infection (Figure 6B).

Altogether, these data establish *CgEP4* as a key virulence factor that promotes fungal entry, colonization, and suppression of host defences. Its absence results in reduced *in planta* growth, delayed penetration, and enhanced papilla-mediated resistance, supporting a role in modulating host responses from the earliest stages of the interaction.

### CgEP4 contributes to fungal development and stress tolerance

Beyond their roles in host colonization, CgEP1 and CgEP4 appear to influence general fungal physiology. An *in vitro* phenotypic assay analysis was conducted on the mutants Δ*CgEP1* and Δ*CgEP4*. Compared with the wild-type strain M1.001, both mutants showed reduced radial growth on potato dextrose agar (PDA), with Δ*CgEP1* exhibiting a stronger defect in colony expansion (Figure S6). Asexual development was also affected: Δ*CgEP1* produced significantly fewer conidia and showed a marked reduction in germination efficiency, while Δ*CgEP4* displayed milder but consistent defects in both traits (Figure S6).

Under abiotic stress conditions, the two effectors showed distinct but overlapping contributions. NaCl exposure increased growth inhibition for both mutants compared to M1.001 while Δ*CgEP1* showed stronger inhibition effects. A similar pattern was observed under potassium chloride (KCl) treatment, where Δ*CgEP1* again exhibited significantly higher inhibition than the other strains, including Δ*CgEP4* and the wild type. These results suggest that CgEP1 plays a prominent role in managing ionic stress, regardless of the specific ion. Upon exposure to hydrogen peroxide (H₂O₂), Δ*CgEP1* also showed heightened sensitivity, while Δ*CgEP4* maintained intermediate growth, indicating that both effectors contribute to oxidative stress resistance, with CgEP1 having a greater impact under these conditions.

We examined cell envelope integrity through tests that evaluated mutant responses to agents that disrupt cell wall and plasmatic membrane components. No significant differences were observed on SDS-containing media for Δ*CgEP4*. However, Δ*CgEP1* showed increased sensitivity to Uvitex 2B, indicating an altered response to chitin-binding stress. Δ*CgEP1* showed a higher sensitivity to SDS stress, suggesting a defect in plasma membrane stability. In contrast, Δ*CgEP4* grew more robustly than the wild type on Congo Red, indicating a moderate increase in resistance to cell wall interference while Δ*CgEP1* had an increased sensitivity to Congo Red (Figure S7).

Collectively, these results demonstrate that both CgEP1 and CgEP4 affect core physiological processes beyond infection. While CgEP1 appears to be more critical for conidiation, germination, and tolerance to ionic and oxidative stress, as well as resistance to structural stress, CgEP4 plays a broader role that includes vegetative growth and resistance to Na^+^ ionic stress. The observed functional diversity following gene duplication indicates that each effector provides unique contributions to environmental fitness in the fungus.

### CgEP4 modulates maize and fungal gene expression during infection

Loss of CgEP4 triggered a focused but consistent shift in the maize transcriptome. Transcriptomic analysis using RNA-seq identified 205 genes whose expression changed in Δ*CgEP4* infections compared with WT (padj < 0.05; |log₂FC| ≥ 2; Figure S8). Of these, 117 genes were uniquely affected in this comparison (Table S3), suggesting that they specifically respond to the presence or absence of the effector. 61 were upregulated in the mutant during infection of the plant, including the transcription factors bZIP-1 (*Zm00001eb157860*) and ARF-TF2 (*Zm00001eb045640*), the metabolic enzymes triose-phosphate isomerase 3 (*Zm00001eb335870*) and starch synthase I (*Zm00001eb376100*), and the cytokinin-degrading enzyme CKO-4 (*Zm00001eb151670*). This cohort yielded strong GO enrichment for “MAPK cascade” and “MAP-kinase activity” (Figure 7). Conversely, 56 genes were downregulated when CgEP4 was absent; this set included the defence enzyme CHI-B1 (*Zm00001eb425600*; log₂FC = -2.7), the immune regulator WRKY-92 (*Zm00001eb350280*), the splicing factor SR45-1 (*Zm00001eb140450*), THO-complex subunit 5 (*Zm00001eb255990*) and the metabolic enzyme IDH-5 (*Zm00001eb183760*), producing GO enrichment for “RNA processing / spliceosome” and “protein processing in endoplasmic reticulum” (Table S3) (Figure 7). Thus, CgEP4 normally suppresses MAPK-linked signalling, hormone and carbohydrate metabolism while sustaining transcripts involved in defence, RNA maturation and proteostasis; its deletion reverses this balance, mirroring the reduced penetration efficiency and fungal biomass of the Δ*CgEP4* strain. Principal component and volcano plots illustrating these global shifts are provided in Figure S9.

**Figure 7.**
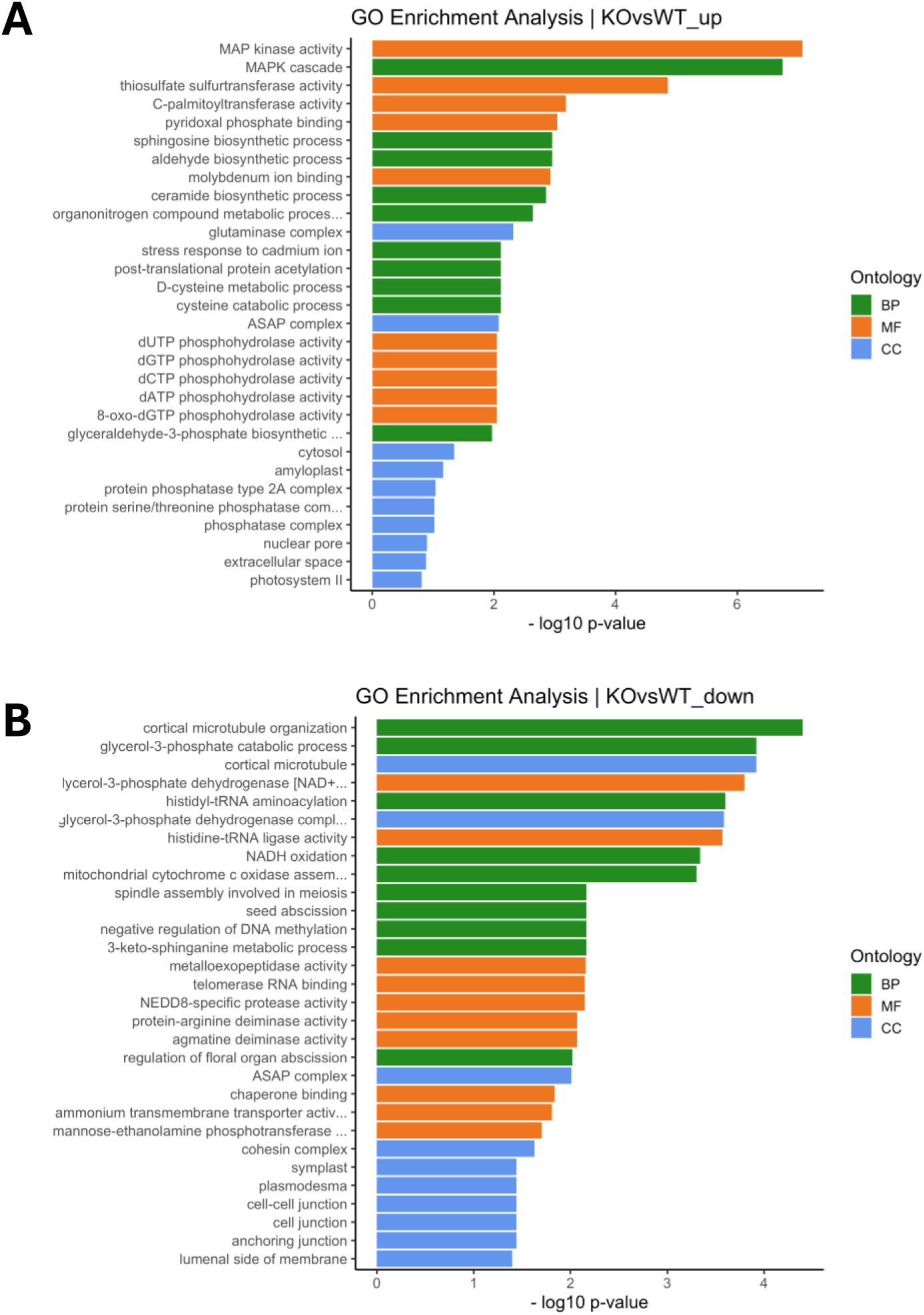
Transcriptomic analysis. **(A)** GO enrichment analysis of *Zea mays* DEGs upregulated in the KO vs. WT comparison. Enriched terms include MAP kinase activity, sulfur metabolism, and stress-related genes. **(B)** GO enrichment analysis of DEGs downregulated in the KO vs. WT comparison. Enriched terms include microtubule organization, redox processes, and metabolic pathways. Colours indicate GO categories: green = Biological Process (BP), orange = Molecular Function (MF), and blue = Cellular Component (CC).

Analysis of *C. graminicola* transcripts in infected leaf tissue identified 219 fungal DEGs that were exclusive to the KO vs. WT comparison (Table S4). Only five genes were upregulated in the mutant: a putative 4-hydroxy-acetophenone mono-oxygenase (*CGRA01v4_10455*, log₂FC = +22.7), a TauD-family taurine dioxygenase (CGRA01*_14901*, +22.6), a haloacid dehalogenase-like hydrolase (CGRA01*_14502*, +12.0), an uncharacterized protein (CGRA01*_07957*, +9.8) and a mitochondrial Mn-superoxide dismutase (CGRA01*_14270*, +3.1). All five encode oxidoreductases or detoxification enzymes, suggesting a compensatory stress response.

By contrast, 216 genes were downregulated, including 38 predicted secreted proteins—among them four LysM effectors, six glycoside-hydrolase CAZymes and three small cysteine-rich proteins—as well as multiple ribosomal proteins, amino-acid biosynthetic enzymes and chitin-synthase regulators. GO enrichment highlighted “ribosome biogenesis” and “cell-wall organization”, indicating that it is required to maintain both the secretory effector load and basic growth machinery during host colonization (Table S4). Principal component and volcano plots illustrating these global shifts are provided in Figure S10.

## Discussion

Our research identified CgEP4, novel effector protein from *C. graminicola* which contributes to both host infection and fungal fitness. CgEP4 is secreted, after which it translocates to host nuclei during early infection to facilitate penetration and disease progression. The absence of CgEP4 led to reduced fungal virulence on maize leaves and stalks together with delayed penetration through epidermal tissue, increased papilla formation and reduced fungal growth inside the plant. In addition, the saprophytic development of the CgEP4 mutant was impaired, characterized by reduced vegetative growth, as well as compromised conidiation and increased sensitivity to oxidative and ionic stress.

Our results show that CgEP4 is a dual-role effector, promoting fungal development and manipulating the host by suppressing early defence responses and targeting maize nuclei. Increasing evidence suggests that a single effector can coordinate fungal development and immune evasion (Feldman et al., 2020). For instance, the metalloprotease effector UmFly1 from *Ustilago maydis* targets both fungal and plant chitinases, promoting fungal cell separation while suppressing host immune responses (Ökmen et al., 2018). These findings underscore that dual-function effectors may be an adaptive strategy among plant-pathogenic fungi

The evolutionary reconstruction of the *CgEP1/CgEP4* gene family highlights a dynamic pattern of duplication, loss, and functional diversification, tightly associated with the ecological and evolutionary history of *Colletotrichum* species. The Oligocene–Miocene duplication event (DE2), which gave rise to CgEP1 and CgEP4, occurred approximately 28.2–17.8 million years ago, a period of profound global environmental change. The transition from the Oligocene to the Miocene marked the early expansion of C4 grasses, which transformed terrestrial ecosystems by promoting the spread of open grasslands, particularly during the late Miocene (Baroncelli et al., 2024). This major shift in plant communities likely reshaped plant–pathogen interactions, creating new ecological niches and selective pressures. In this context, the emergence of CgEP4 may represent an adaptive response to the diversification of grass hosts, potentially contributing to the specialization of *Colletotrichum graminicola* and related species toward monocot hosts. The coexistence of both CgEP1 and CgEP4 within the *Graminicola* complex, coupled with their distinct functional roles, suggests subfunctionalization or neofunctionalization following duplication. The presence of two major clades encompassing nearly all *Colletotrichum* complexes, together with the pattern of single-copy retention in early-diverging lineages such as the *Orbiculare* complex, supports the hypothesis of an ancient duplication (DE1) before the genus diversified. Subsequent lineage-specific losses and duplications (DE2 and DE3) have shaped the present distribution of this gene family. The expansion of this effector family within the Graminicola complex, combined with evidence for functional divergence, underscores its likely contribution to the adaptive evolution of pathogenicity in grass-infecting *Colletotrichum* species. Interestingly, CgEP1 was previously shown to be subject to positive selection, consistent with the arms race hypothesis in which pathogen effectors are constantly evolving in response to evolutionary changes in the components of the host with which it interacts (Vargas et al., 2016). In contrast, CgEP4 is highly conserved with no evidence of positive selection. Thus, while both genes are clearly related by an ancestral gene duplication, their evolutionary paths since that duplication have become distinct. These findings suggest that the diversification of the *CgEP1/CgEP4* family paralleled major ecological transitions in grass evolution, potentially providing new molecular tools for successful colonization of emerging hosts.

The localization of *CgEP4* and *CgEP1* in repeat-rich regions of chromosome 6 is consistent with the compartmentalized genome structure reported in *Colletotrichum*, where effectors and secondary metabolism genes are associated with transposable elements (TEs). In *C. graminicola*, secreted proteins, including effectors, are located near repetitive elements in AT-rich, gene-poor regions (Becerra et al., 2023). Such associations, involving Gypsy-type LTRs, mini-chromosomes, and repeat-driven rearrangements, have been described in *C. truncatum*, *C. higginsianum*, or *C. lindemuthianum* (Da Silva et al., 2024; Dallery et al., 2017; Gan et al., 2013; Rao et al., 2018; Tsushima et al., 2019). These studies support a model in which TE-rich compartments promote both the emergence and diversification of effectors, likely including the ancestral duplication event that gave rise to *CgEP1* and *CgEP4*.

The nuclear localization of CgEP4 during early infection indicates that it regulates host gene expression or chromatin-related processes from inside the nucleus. Our fluorescence microscopy imaging showed consistent accumulation of CgEP4 in host nuclei, where it may act by directly binding to chromatin or host DNA to reprogram gene expression, or by hijacking or destabilizing nuclear regulatory proteins such as transcription factors, co-activators, or repressors, thereby altering transcriptional dynamics to favour colonization (Kim et al., 2020; Yang et al., 2022). This nuclear behaviour reflects that of its paralog CgEP1, which binds to host chromatin to control transcriptional activity and promotes susceptibility (Vargas et al., 2016). The two effectors CgEP4 and CgEP1 might work together through sequential or synergistic mechanisms by targeting separate host nuclear regulators or gene networks. Nuclear targeting appears to be a recurrent strategy among fungal effectors to manipulate host transcriptional or regulatory hubs. The nuclear protein CgNLP1 from *Colletotrichum gloeosporioides* interferes with MYB transcription factor localization to suppress cell death in rubber tree (Yang et al., 2022), while the conserved effector candidate CEC3 triggers nuclear expansion and cell death in *Nicotiana benthamiana*, likely through disruption of nuclear structure or function (Tsushima et al., 2021). Related strategies have also been documented in other filamentous pathogens such as *Magnaporthe oryzae*, *Fusarium oxysporum* and *Verticillium dahliae*, where nuclear effectors influence hormone signalling, chromatin state, or transcriptional control (Chen et al., 2023; Lee et al., 2023; Li et al., 2024a; Yang et al., 2023). These proteins demonstrate the versatility of nuclear effectors across fungal lineages, supporting the idea that CgEP4 may exploit conserved vulnerabilities in host nuclear functions to promote infection. In addition, we observed an apparent enrichment in subnuclear regions morphologically resembling the nucleolus. This pattern suggests that CgEP4 may interact with nucleolar components. Such a localization would be consistent with reports of effectors from other plant pathogens that target the nucleolus to interfere with ribosome biogenesis, RNA processing, or stress-related signalling (Ranty-Roby et al., 2024), but specific nucleolar markers would be needed to assess it.

Given its early expression and nuclear accumulation, CgEP4 is likely to play a key role during the biotrophic phase of infection, potentially delaying host defence activation. The marked attenuation of the virulence of Δ*CgEP4* mutants underscores the importance of this effector for the successful colonization of maize tissues. CgEP4 appears to support both the establishment and development of disease, as evidenced by reduced lesion size and fungal biomass. These observations suggest that CgEP4 functions early in the infection process, helping to establish a compatible interaction while also enabling subsequent fungal proliferation. Its virulence contribution is comparable to other core effectors in *C. graminicola*, such as CgEP1 or the metalloprotease CgFL, reinforcing that CgEP4 plays a non-redundant and central role in pathogenesis (Sanz-Martín et al., 2016a; Vargas et al., 2016).

Deletion of *CgEP4* led to impaired penetration efficiency and a significantly elevated rate of effective papilla formation, demonstrating that this effector plays a central role in suppressing early immune responses at the site of fungal entry. Appressorial penetration in several plant-fungal interactions occurs tightly coordinated with the evasion of localized defence mechanisms that include fast callose and cell wall reinforcement deposition. Similar strategies have been reported in other pathogens: for example, the lipase effector FGL1 from *Fusarium graminearum* releases free fatty acids that inhibit host callose deposition and facilitate tissue colonization (Blümke et al., 2014). In addition, *Puccinia striiformis* effector Hasp155 suppresses chitin-triggered immunity and callose accumulation to promote penetration (Yan et al., 2024). The Δ*CgEP4* strain appressoria frequently failed to penetrate the epidermis, particularly at early time points, which suggests that without this effector, host defences are activated more quickly and effectively, resulting in early entry failure. Although penetration occurs, the infection is delayed, which means that these pre-invasive barriers were not neutralized within a critical time window for penetration to occur.

Beyond their canonical function in manipulating the host, CgEP1 and CgEP4 have diverged to sustain complementary aspects of *C. graminicola* physiology. Deleting either effector compromises vegetative growth and asexual sporulation, yet the spectrum and magnitude of the defects differ: Δ*CgEP1* is most impaired in colony expansion, conidiation and germination, whereas Δ*CgEP4* shows milder reductions but a sharp sensitivity to Na⁺ stress. Sodium toxicity perturbs intracellular ion balance and membrane potential, demanding precise control of ENA-type ATPases and Hog1-mediated pathways (Brown et al., 2014; Hohmann, 2015; Jiang et al., 2018); the nuclear localization of CgEP4 therefore hints that it modulates transcriptional programmes required for sodium homeostasis. Conversely, Δ*CgEP1* is hypersensitive to K⁺, SDS, Congo Red and Uvitex 2B, pointing to a broader failure in cell-envelope integrity and oxidative-stress detoxification—traits commonly governed by AP1-like regulators in other phytopathogens (Montibus et al., 2013; Temme & Tudzynski, 2009). These phenotypic partitions, together with the effectors’ paralogous origin, fit a model of post-duplication neofunctionalization in which CgEP1 has specialized in safeguarding envelope stability and ROS homeostasis, while CgEP4 underpins growth and sodium-specific resilience. Such functional bifurcation likely maximizes fungal fitness in the fluctuating rhizosphere and may partly explain the pronounced loss of virulence observed when either gene is deleted: a host that still mounts ROS bursts or ionic defences cannot be efficiently colonized. Effector diversification therefore extends beyond host subversion to reinforce the pathogen’s own environmental robustness, underscoring the intimate evolutionary coupling of developmental and virulence determinants (Li et al., 2024b; Rokas et al., 2018).

Loss of *CgEP4* triggers a robust MAPK-mediated immune response in maize. Within 48 hours post-inoculation, Δ*CgEP4* plants showed strong induction of canonical defence-related genes, including *CHI-B1*, *WRKY92*, and *bZIP1*, alongside hormonal regulators such as *CKO4* (cytokinin oxidase) and *ARF-TF2* (auxin response factor). This pattern is indicative of an early transcriptional reprogramming aimed at cell wall fortification and immune signalling, matching the increased papillae deposition and delayed penetration observed cytologically. This host response resembles those observed during infection with nuclear-localized effectors such as MoHTR3 from *Magnaporthe oryzae*, which modulates the expression of defence-associated hormone signalling genes via chromatin interaction in the nucleus (Lee et al., 2023), and ArPEC25 from *Ascochyta rabiei*, which interferes with host transcription factors to suppress the phenylpropanoid pathway and reduce lignin biosynthesis (Singh et al., 2023).

Interestingly, maize genes involved in RNA processing (*SR45-1*, *THO5*) and central metabolism (*IDH5*, glycolytic and starch synthesis enzymes) were significantly downregulated in Δ*CgEP4*-infected tissues. This suggests that CgEP4 not only suppresses defence gene activation but also safeguards host biosynthetic capacity during biotrophic colonization. Such a dual function aligns with the nuclear localization of CgEP4 and parallels the behaviour of nuclear effectors like *CgEP1* or *MoHTR3*, which maintain host cell viability while suppressing immune activation.

On the fungal side, transcriptomic analysis of Δ*CgEP4* revealed widespread repression of core pathogenicity pathways. Expression of 216 fungal genes was reduced, including 38 encoding predicted effectors, multiple ribosomal proteins, and key regulators of chitin biosynthesis. In contrast, only a few genes involved in detoxification were upregulated. This expression profile suggests that CgEP4 helps maintain fungal biosynthetic and secretory activity under immunologically hostile conditions. Once host defences are activated, the fungus appears to downregulate virulence and growth-associated genes, possibly as a stress response or survival strategy.

Taken together, these dual RNA-seq results underscore the role of CgEP4 as a cross-kingdom effector: by dampening host defence signalling and stabilizing host metabolic outputs, it establishes a permissive environment; simultaneously, it sustains the fungal gene expression machinery required for colonization. In its absence, both the host and pathogen undergo transcriptional changes, demonstrating how a single nuclear effector can synchronize immune suppression and biotrophic expansion to facilitate successful infection.

### Experimental procedures

#### Strains and growth conditions

*Colletotrichum graminicola* wild-type strain M1.001 (Forgey et al., 1978) and the corresponding mutants and complemented strains were routinely cultured on potato dextrose agar (PDA; Difco, Becton, Dickinson and Company, Franklin Lakes, NJ, USA) at 23 °C under continuous white light. For liquid cultures, potato dextrose broth (PDB; Difco, Becton, Dickinson and Company, Franklin Lakes, NJ, USA) was used to produce vegetative mycelia for DNA or RNA extraction. Conidial production was induced on PDA plates incubated for 7 days, and conidia were harvested by flooding plates with sterile distilled water and filtering through double-layered cheesecloth. All strains were stored in silica stocks at −80 °C for long-term preservation.

#### Plant cultivation

Maize plants of the derived inbred line Mo940 (Warren, 1977) were cultivated until they reached the stage of development V3 (approximately 3 weeks old) at 25 °C under long-day conditions (16 h of light and 8 h of dark) and 80% humidity (Vargas et al., 2012).

#### Gene structure validation

To verify the structure of *CgEP4,* total RNA was extracted from infected maize leaves at 48 hpi and reverse transcribed into cDNA. RNA was isolated using the RNeasy Plant Mini Kit for RNA Extraction (Qiagen, Hilden, Germany), and cDNA was synthesized from 800 ng of RNA using the PrimeScript RT Reagent Kit (Takara Bio Inc., Shiga, Japan). A region spanning the predicted intron was amplified by PCR using flanking primers (Table S5), cloned into a pCR™8/GW/TOPO®TA (Invitrogen, Thermo Fisher Scientific), and sequenced. The resulting cDNA sequence was aligned to the reference genomic model using Geneious Prime 2025.0.3 (https://www.geneious.com) to confirm intron length.

#### Modelling of 3D structure

The modelling of the 3D structure was performed using AlphaFold2 (AF), which is included in the ColabFold pipeline, employing the pdb100 option as the template_mode to detect structural template homologues similar to CgEP4 (Jumper et al., 2021; Mirdita et al., 2022). The modelling was constructed with an MSA generated to infer a hypothetical three-dimensional structure of CgEP4. *Ab initio* structural modelling was performed with MMseqs2 and HHsearch as implemented in ColabFold (Fidler et al., 2016; Mirdita et al., 2022). The refinement of the best model was performed based on the per-residue confidence score (plDDT) suggested by AF. Model preparation for molecular dynamics (MD) was performed using GROMACS version 2022 (Abraham et al., 2015). The protein setup was conducted by using SPC/E as water model and the OPLS/AA Force Field. The protein was solvated in a water box with 1.0 angstrom of length per side and an isotonic concentration of NaCl. Energy minimization, NPT and NVT equilibration were conducted with a trajectory of 100,000 steps. MD simulation was performed in an NPT environment during 2 ns. The RMSD trajectory was visualized by using the Bio3D Package in R (Grant et al., 2006). The 3D structure of CgEP4 was visualized in Discovery Studio Client (BIOVIA DS, 2017).

#### Phylogenetic analysis

The evolutionary relationships of CgEP1 and CgEP4 were investigated through comparative phylogenomic analyses across 40 publicly available *Colletotrichum* reference proteomes (Table S6). The CgEP1 protein sequence was initially used as a query in a BLASTp search against all these proteomes to identify putative homologs. Homologous sequences identified in this search were used as query sequences for a second round of BLASTp searches. This process was continued iteratively until no new homologous sequences were identified. A total of 60 protein sequences showing significant similarity (E-value < 1e−5) were retrieved. Multiple sequence alignment was performed using MAFFT v7 (Katoh & Standley, 2013) with default parameters. Maximum likelihood phylogenetic inference was carried out using PhyML (Guindon et al., 2010) under the LG+G+I model, with branch support estimated from 1,000 bootstrap replicates. The resulting tree was visualized and manually inspected in Geneious Prime 2025.0.3 (Biomatters Ltd., 2025).

To contextualize effector diversification within the broader evolutionary framework of the genus, a *Colletotrichum* species tree was also inferred. Orthologous genes were identified across the same 40 proteomes using OrthoFinder v2.5.5 (Emms & Kelly, 2019), employing default inflation parameters. Single-copy orthogroups were extracted, and the corresponding protein sequences were individually aligned with MAFFT, concatenated into a supermatrix, and used to infer a species-level phylogeny with FastTree v2.1.11 (Baroncelli et al., 2024; Price et al., 2010). The resulting tree topology was consistent with previously published *Colletotrichum* phylogenies and served as a reference for interpreting the diversification patterns of CgEP1 and CgEP4 within the genus.

#### Subcellular localization

Nuclear localization of CgEP4 was assessed using both *in silico* predictions and *in planta* fluorescence imaging. Nuclear localization signals (NLSs) were predicted using multiple online tools (Table S2), with and without the signal peptide sequence. Translational fusions were constructed between the *CgEP4* coding sequence (with or without the signal peptide) and mCherry, driven by the native *CgEP4* promoter (Figure S3). The stop codon was removed to generate C-terminal fusions, and constructs were cloned into the pFPL-Rh vector via Gateway cloning following the method of Gong et al., (2015). In addition, a constitutive eGFP fusion lacking the signal peptide (*PgpdA::CgEP4^ΔSP^–eGFP*) was generated using the same pFPL cloning system but with the pFPL-Gh vector and the *A. nidulans gpdA* promoter instead of the native *CgEP4* promoter (Figure S4). Constructs were transformed into *C. graminicola* M1.001 protoplasts and verified by PCR (Sukno et al., 2008; Thon et al., 2002). Transformed strains and wild type were used to infect maize leaf sheaths. Infected tissues were collected at 36 hpi, stained with DAPI, and visualized using a DMi8 inverted fluorescence microscope (Leica Microsystems, Wetzlar, Germany). Images were processed and analysed using Fiji/ImageJ (Schindelin et al., 2012).

#### Gene deletion and complementation

The *CgEP4* gene was deleted by homologous recombination using the Double-Joint PCR method (Yu et al., 2004). Flanking regions (5′ and 3′ UTRs) of *CgEP4* were amplified with primer pairs listed in Table S5 and fused to a hygromycin resistance cassette. The resulting construct was introduced into *C. graminicola* M1.001 protoplasts by PEG-mediated transformation, as previously described (Thon et al., 2002; Sukno et al., 2008) (Figure S11). Putative Δ*CgEP4* transformants were screened by PCR and further confirmed by Southern blot analysis using DIG-labelled probes specific to the 5′ region and *hph* gene. For complementation, the *CgEP4* coding sequence with its native promoter was cloned into the pKW4 vector and reintroduced into the Δ*CgEP4* background. Positive transformants were confirmed by PCR and number of copies and correct integration in the genome was checked by Southern blot using DIG-labelled probes specific to the 5′ region and *nat* gene (Figure S12).

#### Expression profile

Expression of *CgEP4* during maize infection was analysed by quantitative real-time PCR (qRT-PCR). RNA was extracted from *in vitro*-grown mycelium and conidia, as well as from infected maize leaves collected at 24, 36, 48, 60, and 72 hours post-inoculation (hpi), and at 4 and 8 days post-inoculation (dpi), following the protocol of Vargas et al. (2012). RNA was isolated using the RNeasy Plant Mini Kit for RNA Extraction (Qiagen, Hilden, Germany), and cDNA was synthesized from 800 ng of RNA using the PrimeScript RT Reagent Kit (Takara Bio Inc., Shiga, Japan). Reactions were performed with KAPA SYBR Green qPCR Mix (KAPA Biosystems, Wilmington, MA, USA) on a StepOnePlus™ Real-Time PCR System (Applied Biosystems, Foster City, CA, USA) using the following cycling conditions: 95 °C for 5 min; 40 cycles of 95 °C for 10 s, 65 °C for 20 s, and 72 °C for 20 s. Melting curve analysis was included to ensure amplification specificity. Expression levels were normalized to *C. graminicola* histone H3 (Krijger et al., 2008) and relative transcript abundance was calculated using the 2^-ΔΔCt^ method (Livak & Schmittgen, 2001) Primers are listed in Table S5.

#### Anthracnose virulence assays

For leaf blight assays, maize plants (line Mo940) at the V3 stage were laid horizontally on moist trays. Protocol described in Vargas et al. (2012) was followed. Lesion development was evaluated at 3 dpi. For stalk rot assays, 6-week-old maize plants were inoculated at the first internode by pipetting 10 μL of a 1 × 10⁶ spores/mL suspension into a 2-3 mm wound made with a sterile pipette tip (Thon et al., 2000). Plants were incubated in greenhouse conditions (25 °C ± 5 °C, 45% humidity), and stalks were longitudinally split to assess necrotic lesion size at 5 dpi. The lesion size in both assays was measured using Paint.net v5.0.6 (dotPDN LCC).

#### Quantification of fungal biomass

Fungal biomass in infected maize tissue was quantified by qPCR at 3 dpi. Lesions were sampled and total genomic DNA was extracted using the CTAB protocol (Irfan et al., 2013). Quantitative PCR was performed using 10 ng of genomic DNA as template and the KAPA SYBR Green qPCR Mix (KAPA Biosystems). Reactions were carried out on a StepOnePlus™ Real-Time PCR System (Applied Biosystems) with the same cycling conditions described above. Relative fungal biomass was calculated using the 2^^–ΔΔCt^ method (Livak & Schmittgen, 2001), comparing the abundance of the *C. graminicola* ITS2 region to that of maize elongation factor 1 alpha (ZmEF1α) (Lin et al., 2014; Weihmann et al., 2016). Primers used are listed in Table S5.

#### Epidermis penetration and papillae apposition assays

Maize leaves were inoculated with conidia of *C. graminicola* wild type and Δ*CgEP4* strains at a concentration of 1 × 10⁵ spores/mL following Vargas et al. (2012). Leaf sheaths were harvested at 36 and 48 hpi for microscopic analysis. Samples were fixed and bleached in ethanol:glacial acetic acid (3:1, v/v) for 24 h, rehydrated in distilled water, and stored in lactoglycerol (1:1:1, lactic acid:glycerol:water). For staining, tissues were incubated in 0.05% toluidine blue O (in 0.05 M citrate buffer, pH 3.5) for 25 h at 4 °C in the dark to reduce autofluorescence. After sequential washes in distilled water and 0.07 M K₂HPO₄ (pH 8.9), samples were stained with 0.01% aniline blue (in 0.07 M K₂HPO₄) for ≥2 h in the dark This method is an adaptation of different protocols described in Fernandez & Heath (1986) and Stadnik & Buchenauer (2000). Fungal penetration and papilla formation were observed under brightfield and epifluorescence microscopy. The percentage of successful penetration and effective papillae was scored from multiple infection sites per replicate.

#### Abiotic Stress and Cell Wall Integrity Assays

Δ*CgEP1* and Δ*CgEP4* mutants, along with the wild-type and complemented strains, were tested for sensitivity to abiotic and cell wall stressors. Radial growth was assessed on PDA plates supplemented with 0.5 M NaCl, 0.5 M KCl, 0.5 mM H₂O₂, 350 μg/mL Congo Red, 800 μg/mL Uvitex 2B, and 0.01% SDS (Eisermann et al., 2020). Plates were incubated at 23 °C under continuous light, and colony diameters were measured at 7 dpi. Growth inhibition was calculated relative to non-supplemented PDA controls. Each condition was tested in triplicate per strain. Enhanced sensitivity was interpreted as a statistically significant reduction in growth compared to the wild type, while increased tolerance was indicated by reduced inhibition.

#### Transcriptomic analysis

Maize leaves were infected with wild type *C. graminicola*, the Δ*CgEP4* mutant, or mock-inoculated, and tissue was collected at 48 hpi. Three independent biological replicates per condition were used. Total RNA was extracted and sequenced on an Illumina HiSeq 2500 platform (paired-end, 100 bp). Raw reads were quality-checked using FastQC and trimmed with Trimmomatic (Bolger et al., 2014). Cleaned reads were aligned to the *Zea mays* B73_RefGen_v5 genome (Hufford et al., 2021) and the *C. graminicola* CGRA01_v4 genome (Becerra et al., 2023) using HISAT2 (Kim et al., 2019). BAM/SAM files were processed with SAMtools (Li et al., 2009), and transcript abundance was quantified using StringTie-eB (Pertea et al., 2015). Differential expression analysis was performed using DESeq2 (Love et al., 2014). Genes with log₂ fold change ≥ 2 or ≤ –2 and adjusted p-value < 0.05 were considered differentially expressed. Functional annotation of differentially expressed genes was conducted using KEGG (via BLAST-Koala) (Kanehisa et al., 2016), Gene Ontology (Ashburner et al., 2000), and Phytozome (Goodstein et al., 2012) databases. Principal component analyses (PCA) were generated in Python and heatmaps, Venn diagrams and volcano plots were generated in R.

## Acknowledgements

The authors would like to thank the technical assistance of Angélica M. Robledo and Tomás Velasco Criado.

## Supplementary materials

**Figure S1.**
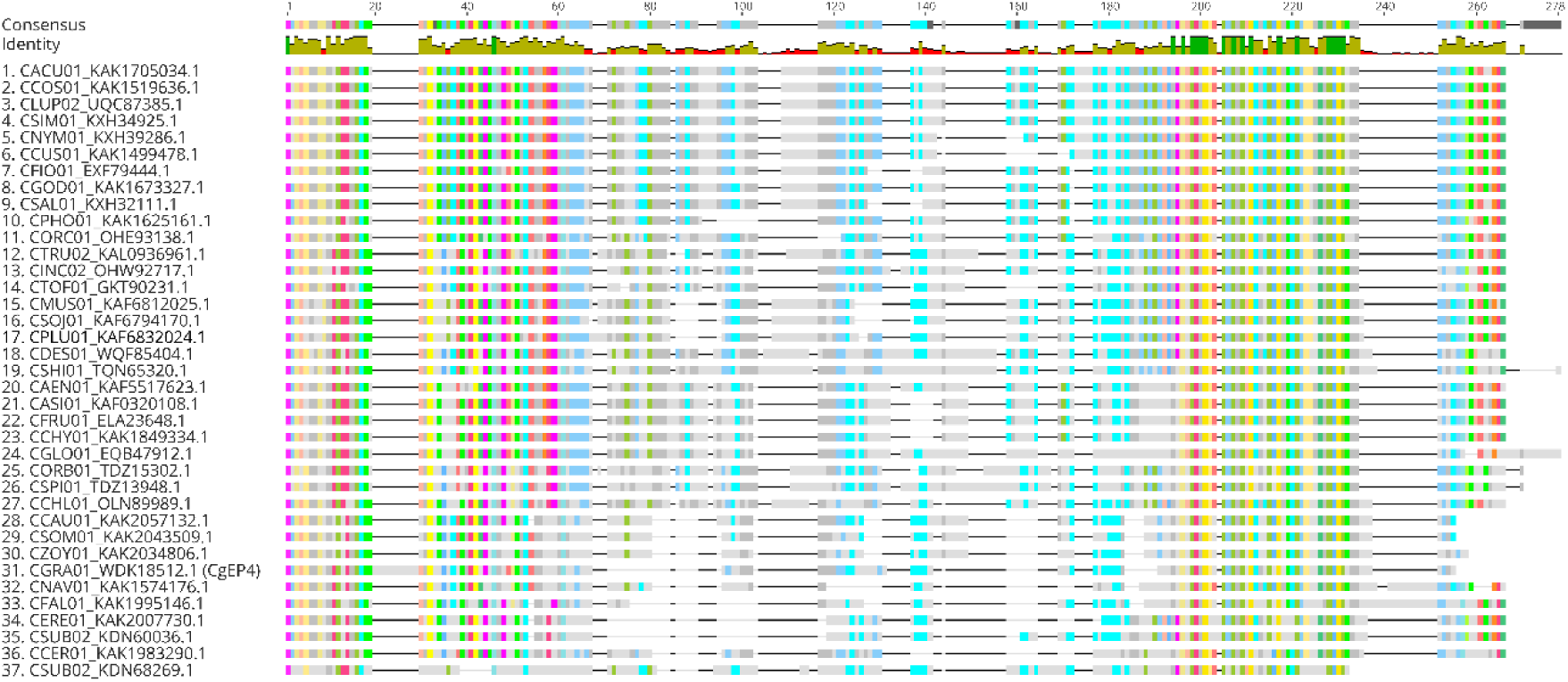
Multiple sequence alignment of CgEP4 orthologs from 36 *Colletotrichum* species. Protein sequences homologous to CgEP4 were retrieved from 41 *Colletotrichum* proteomes using a BLASTp search and aligned with MAFFT v7 (Katoh & Standley, 2013). Only the CgEP4-like sequences from Clade 1 were included; CgEP1-like subclade was excluded for this alignment. The alignment revealed three distinct regions: a conserved N-terminal segment (∼residues 1–60), a highly variable central region (∼residues 61–200), and a conserved C-terminal segment (∼residues 200–end). Coloured blocks indicate amino acid conservation across sequences, while black/grey regions reflect divergent regions with gaps or amino acid substitutions. The consensus and identity plots at the top illustrate residue conservation across the alignment: peaks correspond to highly conserved positions. CgEP4 from *C. graminicola* (WDK18512.1) is included as a reference. Overall, orthologs were found in 36 of 41 genomes analyzed, with pairwise identity exceeding 75 % in the terminal regions but dropping below 40 % in the central domain. The alignment visualization were performed with Geneious Prime 2025.0.3 (Biomatters Ltd., 2025).

**Figure S2.**
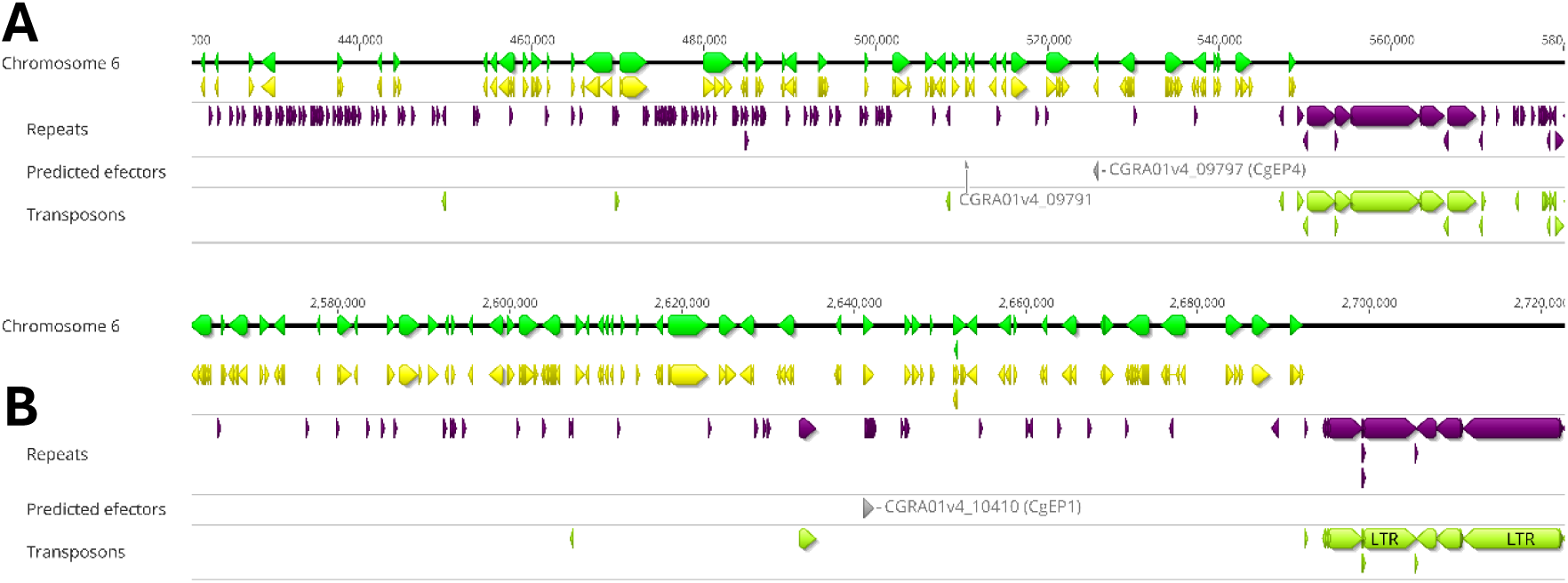
Genomic regions of *CgEP4* and *CgEP1*. **(A)** Genomic landscape surrounding *CgEP4* (*CGRA01v4_09797*) on chromosome 6. The gene is located within a repeat-enriched region and lies adjacent to a full-length LTR retrotransposon. **(B)** Genomic region of *CgEP1* (*CGRA01v4_10410*), also on chromosome 6, similarly positioned in a repeat-rich area with a nearby LTR element. In each panel the tracks, plotted from top to bottom, show protein-coding genes (green arrows), their CDS regions (yellow), annotated repeats (purple), predicted secreted effectors (grey) and full-length transposons (light green). Horizontal rulers show chromosome coordinates (bp). Tracks were represented using Geneious Prime 2025.0.3 (Biomatters Ltd., 2025) based on the genome annotation of Becerra et al. (2023).

**Figure S3.**
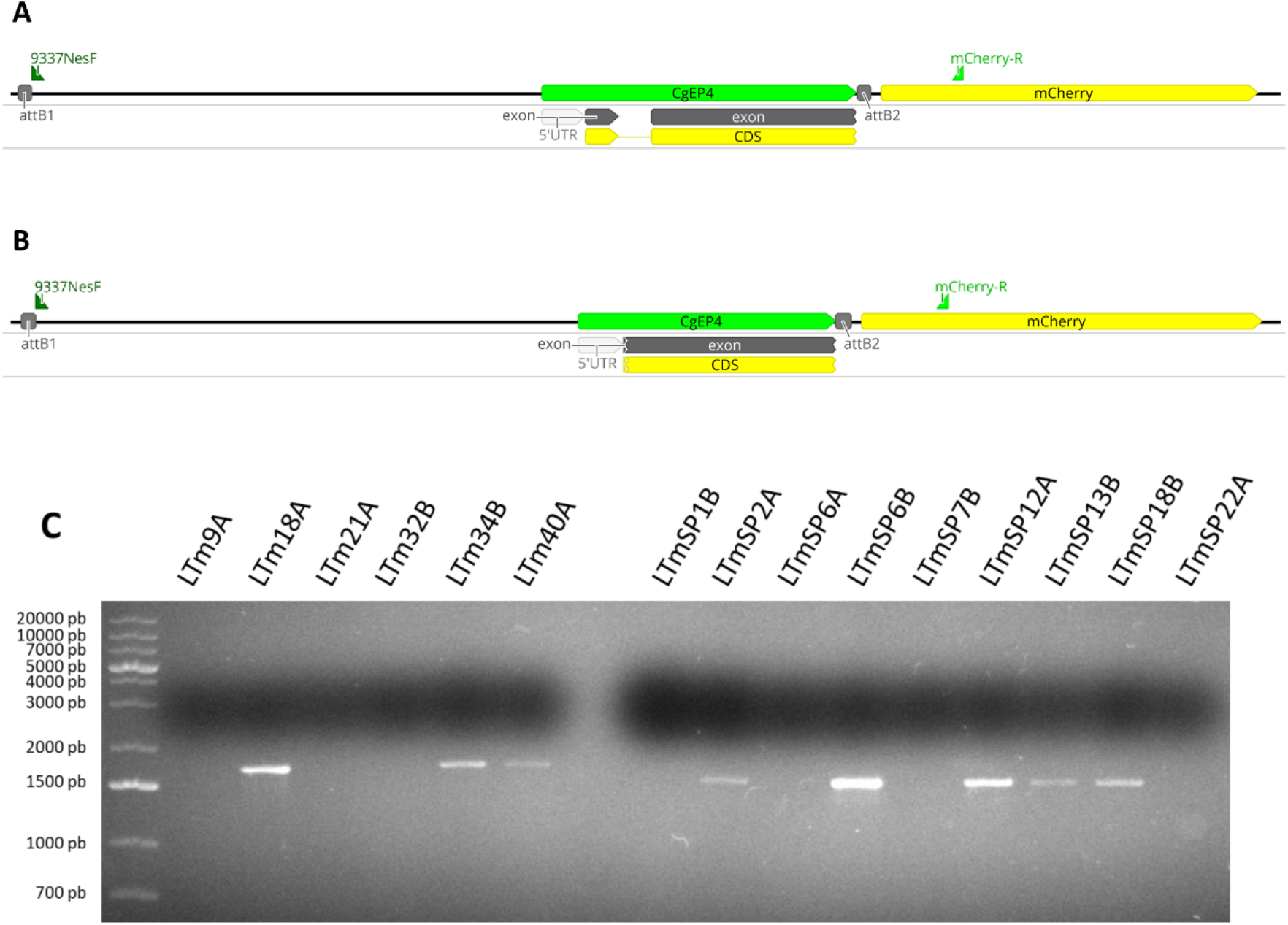
Construction of *CgEP4*-*mCherry* and *CgEP4^ΔSP^*-*mCherry*. **(A)** Schematic diagram of the construct used to investigate the subcellular localization of CgEP4. The native promoter (5′ flanking region) and the full-length coding sequence of *CgEP4* without the stop codon were cloned into the pFPL-Rh vector (Gong et al., 2015), which contains a C-terminal linker*-mCherry* fusion. This allows expression of an in-frame *CgEP4*-*mCherry* fusion protein under native regulatory control. **(B)** The *CgEP4^ΔSP^*-*mCherry* construct was generated in the same way, using the *CgEP4* coding sequence lacking the signal peptide, also cloned without stop codon and fused in-frame to the C-terminal linker*-mCherry* present in the pFPL-Rh vector (Gong et al., 2015). Both constructs were transformed into *C. graminicola* to assess subcellular localization by fluorescence microscopy. **(C)** Representative PCR validation of transformants using primers 9337NesF (forward, located in the 5′ region of the insert) and mCherry Rv (reverse, located within the *mCherry* coding sequence). This primer combination allows confirmation of the correct insertion and in-frame fusion of the *CgEP4* variants. The expected amplicon size is 1699 bp for the full-length construct (*CgEP4-mCherry*) and 1567 bp for the construct lacking the signal peptide (*CgEP4^ΔSP^*-*mCherry*). Transformants LTm18A and LTm34B were selected for further analysis, while LTm6B and LTm18B were used as controls.

**Figure S4.**
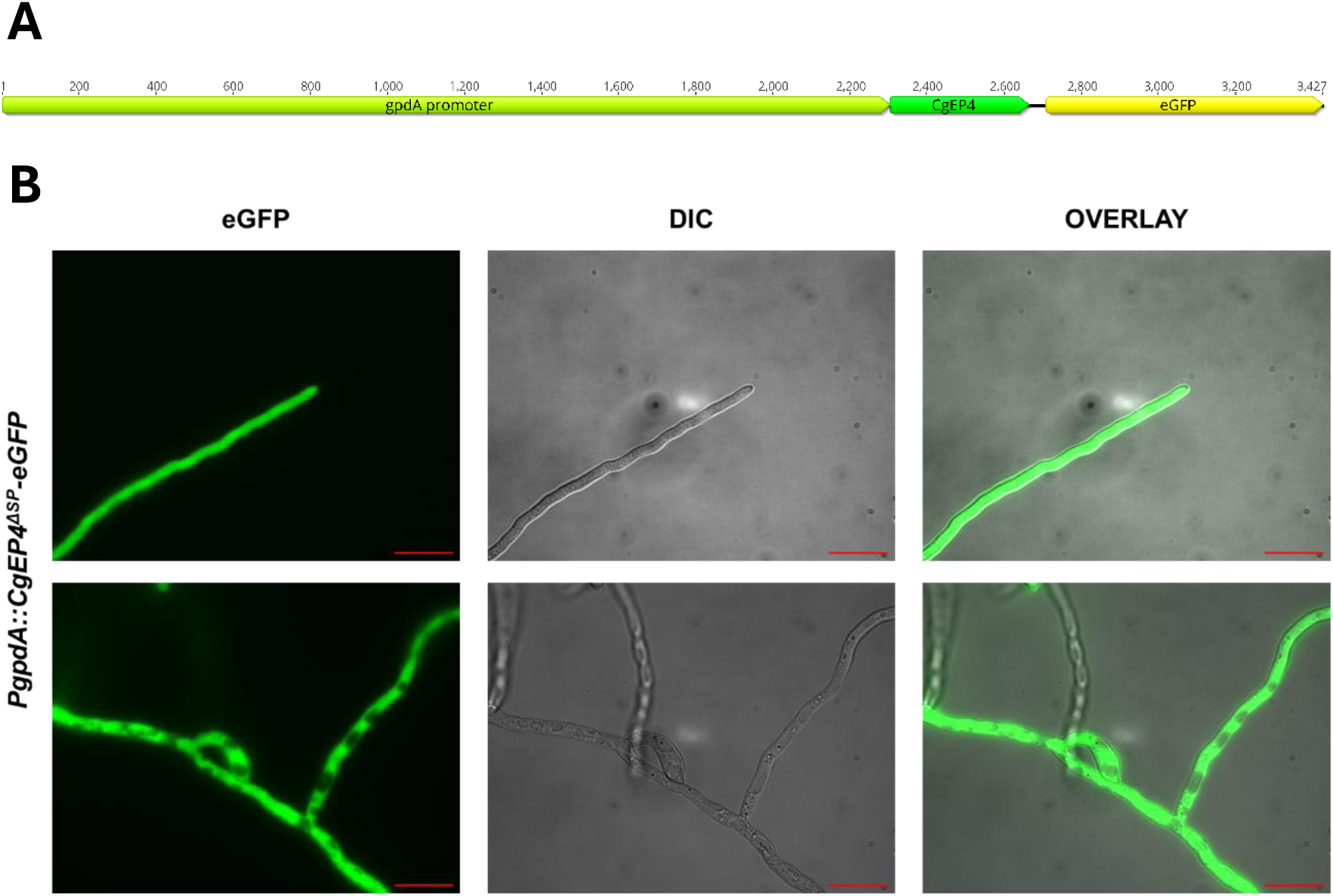
**(A)** Schematic representation of the *PgpdA::CgEP4^ΔSP^–eGFP* construct. The *Aspergillus nidulans gpdA* promoter (light green) drives constitutive expression of *CgEP4* (green) lacking its N-terminal signal peptide. The *eGFP* coding sequence (yellow) was fused in frame to the *CgEP4^ΔSP^* coding sequence via a short linker (black line). **(B)** Fluorescence microscopy of *C. graminicola* hyphae expressing *PgpdA::CgEP4^ΔSP^–eGFP* grown in PDB medium. Strong green fluorescence was observed throughout the cytoplasm of vegetative hyphae, confirming intracellular accumulation of the fusion protein. The absence of fluorescence outside the hyphae indicates that removal of the signal peptide prevents secretion. Scale bars = 20 µm.

**Figure S5.**
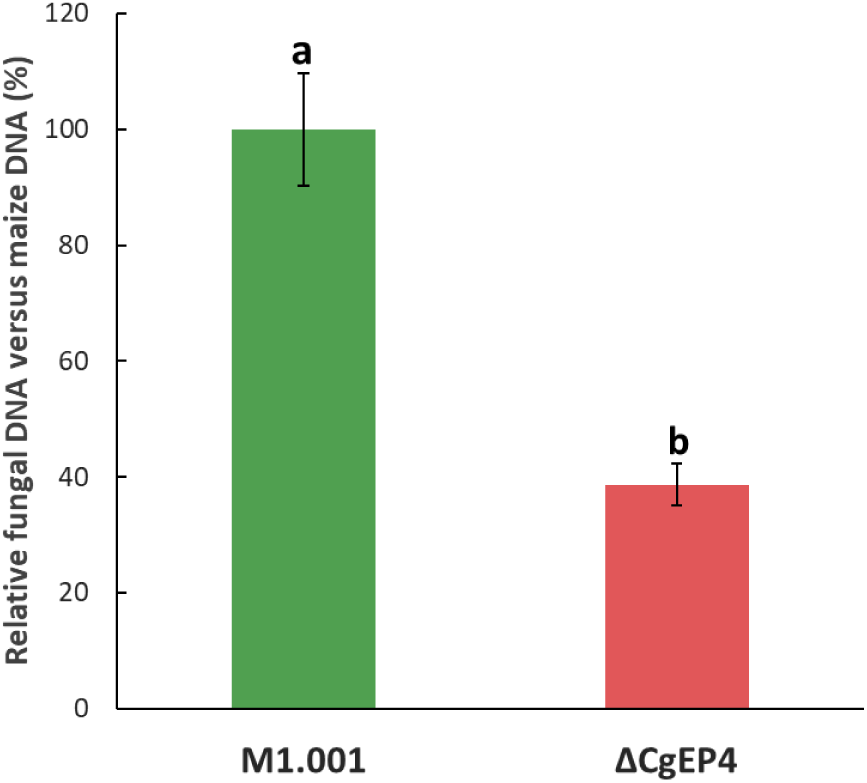
Quantification of fungal biomass in maize leaves at 3 days post-inoculation (dpi). Relative fungal biomass was measured by qPCR in tissues infected with *C. graminicola* wild-type (M1.001) and Δ*CgEP4* strains. Fungal DNA was quantified relative to maize DNA using the 2^⁻ΔΔCt^ method, targeting the *C. graminicola* ITS2 region and maize EF1α as reference genes. Bars represent mean ± standard deviation from three independent biological replicates. Letters denote statistically significant differences compared to the wild type (Student’s t-test, *p* < 0.05). Reduced fungal biomass in Δ*CgEP4* strain infection confirms that the virulence defect is associated with compromised colonization.

**Figure S6.**
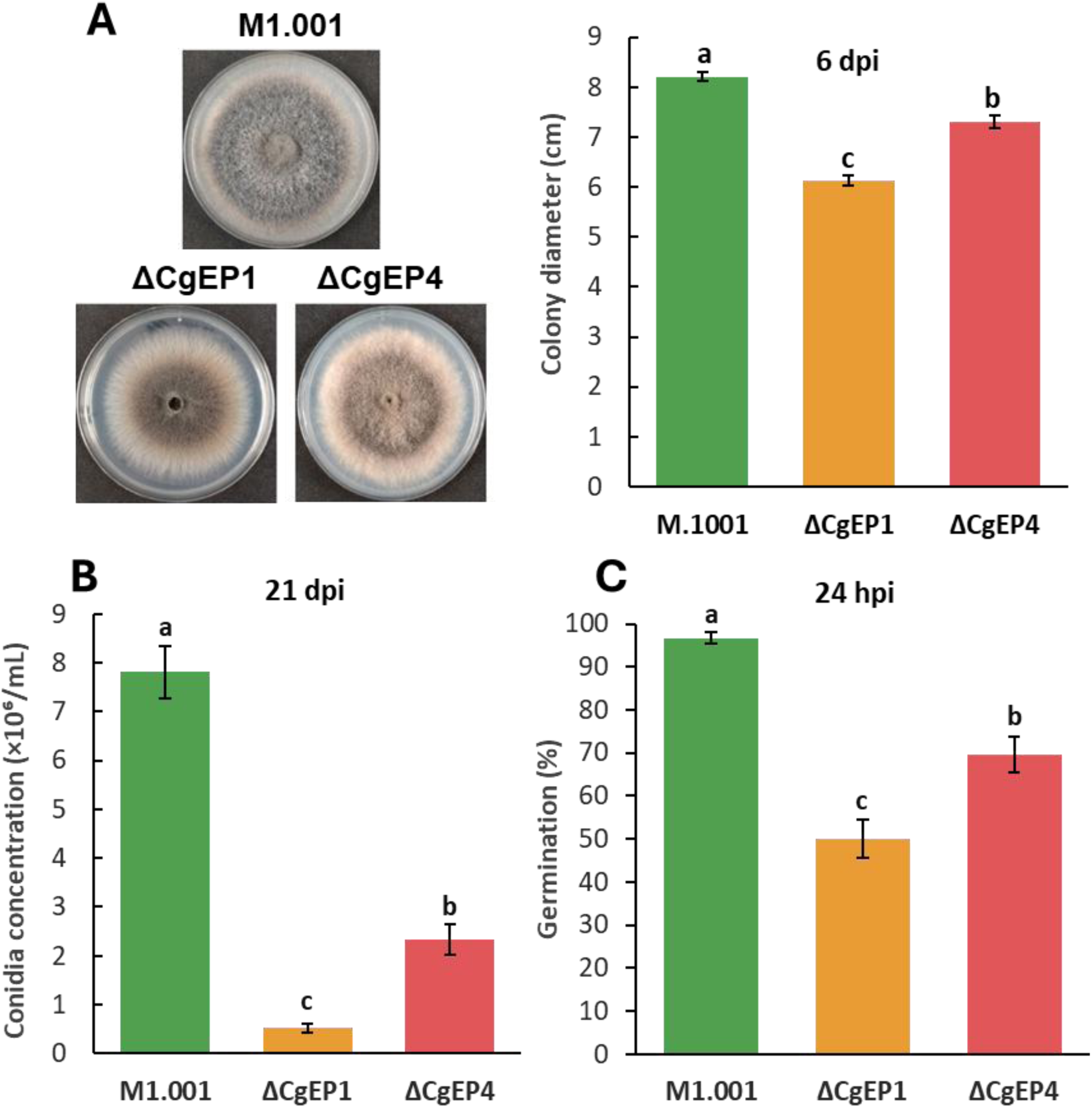
Phenotypic analysis of *CgEP4 and CgEP1* deletion mutants. **(A)** Colony morphology and colony diameter of the wild-type strain M1.001 (WT), Δ*CgEP1*, and Δ*CgEP4* strains on potato dextrose agar (PDA) after six days of incubation at 23 °C under continuous light. Data are from four biological replicates, each comprising at least three technical replicates. **(B)** Conidiation quantified at 21 dpi by counting total spores per Petri dish. The data are expressed as conidia (×10⁶) per millilitre. **(C)** Germination efficiency assessed after 24 hours of incubation on water agar. Bars represent mean values ± standard deviation from three biological replicates with three technical replicates each. Statistical significance was determined using Student’s t-test for all three experiments; different letters indicate significant differences relative to WT (p < 0.05).

**Figure S7.**
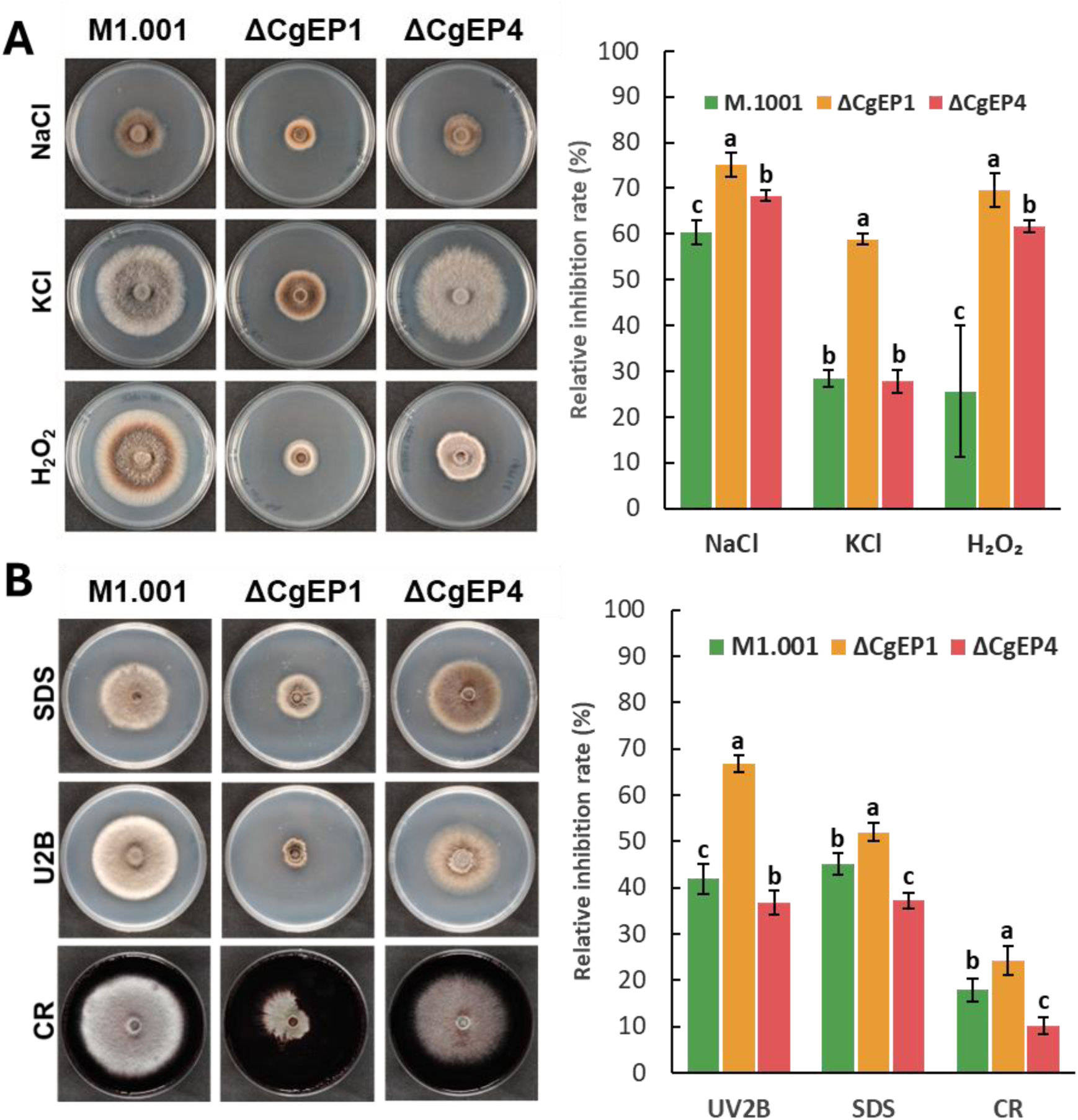
Growth of Δ*CgEP4* under stress conditions. **(A) *Left.*** Colony morphology and growth of WT (*M1.001*), Δ*CgEP1*, and Δ*CgEP4* strains on PDA plates supplemented with osmotic and oxidative stress agents: NaCl (0.5 M), KCl (0.5 M), and H₂O₂ (0.5 mM). ***Right.*** Relative inhibition rate (%) of colony growth under each condition. Inhibition was calculated as [(colony diameter on PDA − diameter under stress) / diameter on PDA] × 100. Bars represent means ± standard deviation from three independent biological replicates. Different letters indicate statistically significant differences between strains under the same treatment (*p* < 0.05, ANOVA followed by Tukey’s test). **(B) *Left.*** Colony morphology and growth of the same strains on PDA supplemented with cell wall–targeting agents: Congo Red (350 µg/mL), SDS (0.01%), and Uvitex 2B (800 µg/mL). ***Right.*** Relative inhibition rate calculated as in (A). Bars represent means ± standard deviation from three independent replicates. Statistical significance was assessed as above.

**Figure S8.**
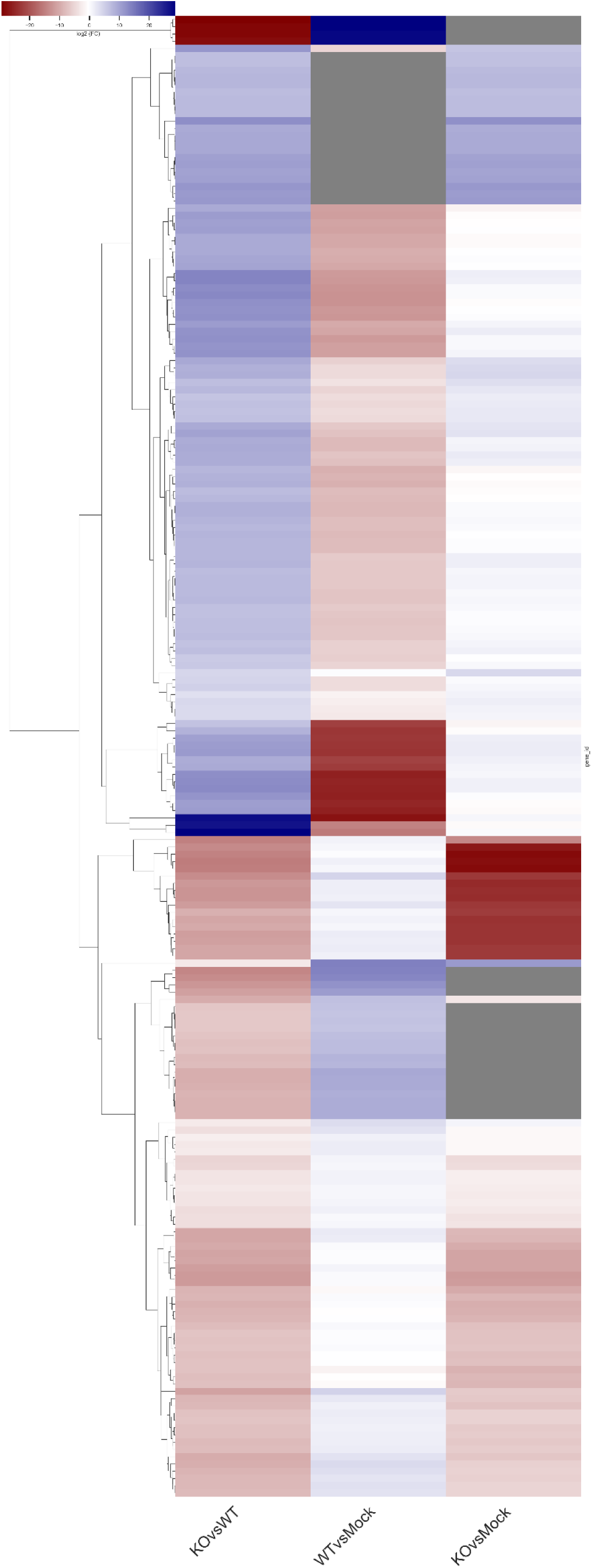
Heatmap of log_2_ fold change (log₂FC) values for all maize genes that were significantly differentially expressed in the comparison KO vs WT. The same genes are shown in the reference contrasts WT vs Mock and KO vs Mock to visualize basal infection responses. Each row represents one gene (n = 205); rows are clustered by complete linkage on distances (dendrogram at left). Colours indicate expression direction and magnitude (scale bar, top): deep red = strong downregulation, white = no change, deep blue = strong upregulation. Grey cells denote cases where no fold change could be calculated because one or both conditions yielded zero mapped reads. The heatmap was generated in Python 3.12.7 using the libraries seaborn 0.13.2 (Waskom, 2021), matplotlib 3.9.3 (Hunter, 2007), pandas 2.3.0 (McKinney, 2010), and scipy 1.13.1 (Virtanen et al., 2020). The complete gene list and differential expression statistics are provided in Supplementary Table S3.

**Figure S9.**
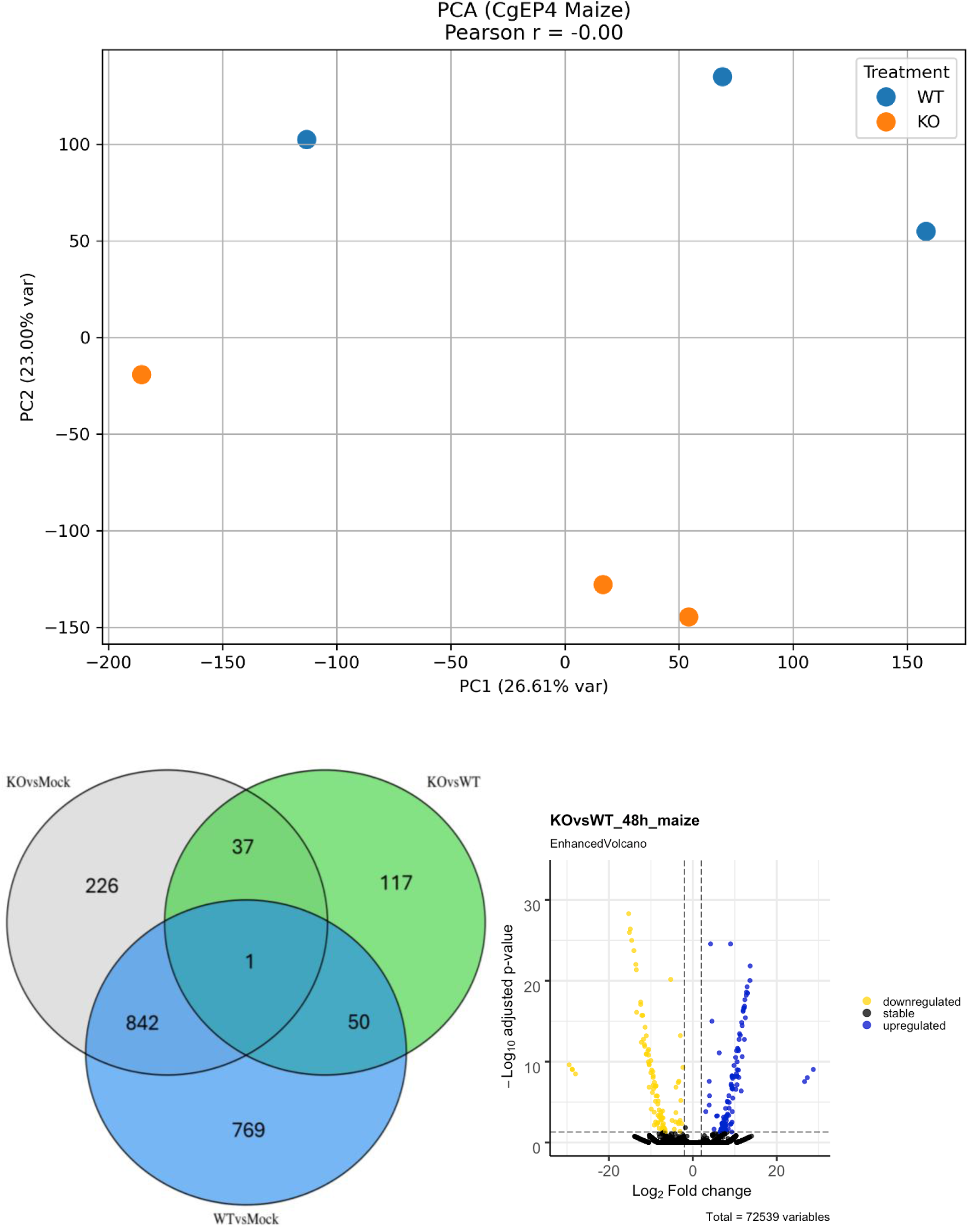
Transcriptomic changes in maize during infection with Δ*CgEP4* mutant strain. **A)** Principal component analysis (PCA) of variance-stabilized read counts for *Zea mays* genes obtained from leaves infected with two strains: M1.001 (WT, blue) and Δ*CgEP4* (KO, orange). Each dot represents an independent biological replicate (*n* = 3 per condition). PCA was performed in Python 3.12.7 (Anaconda distribution on macOS) using pandas 2.3.0 (McKinney, 2010), numpy 2.0.2 (Harris et al., 2020), scikit-learn 1.6.1 (Buitinck et al., 2013), matplotlib 3.9.3 (Hunter, 2007), seaborn 0.13.2 (Waskom, 2021), and scipy 1.13.1 (Virtanen et al., 2020). As input, a gene expression matrix transformed using the variance-stabilizing transformation (VST) from DESeq2 (Love et al., 2014) was used, with grouping by treatment condition. The clustering of replicates within each group indicates the reproducibility of the RNA-seq experiment**. (B)** Venn diagram of differentially expressed genes (DEGs) identified in maize at 48 hpi under three comparisons: KO vs. WT, KO vs. mock, and WT vs. mock. Only DEGs unique to the KO vs WT comparison were used for downstream analysis. Venn diagrams were generated using Intervene (Khan & Mathelier, 2017). **(C)** Volcano plot showing DEGs in maize leaves infected with the Δ*CgEP4* strain relative to the wild type. Genes significantly upregulated (blue) and downregulated (yellow) are shown (|log₂FC| > 2, adjusted *p* < 0.05), as visualized with the EnhancedVolcano package (Blighe et al., 2018). All plots were generated in R 4.0.2 on macOS.

**Figure S10.**
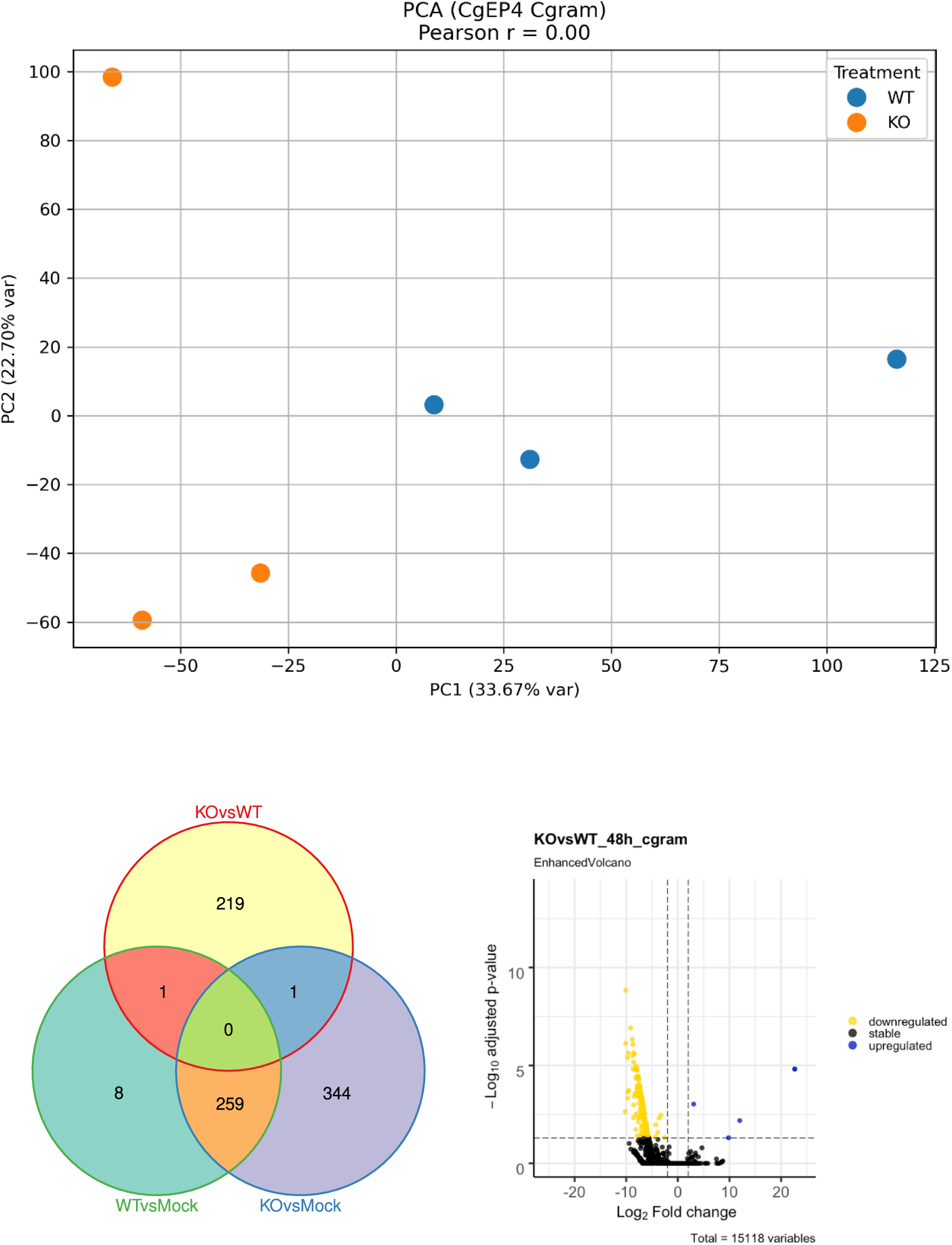
Transcriptomic changes in *C. graminicola* during infection with Δ*CgEP4* mutant strain. **(A)** Principal component analysis (PCA) of variance-stabilized read counts for *C. graminicola* genes obtained from leaves infected with two strains: M1.001 (WT, blue) and Δ*CgEP4* (KO, orange). Each dot represents an independent biological replicate (*n* = 3 per condition). PCA was performed in Python 3.12.7 (Anaconda distribution on macOS) using pandas 2.3.0 (McKinney, 2010), numpy 2.0.2 (Harris et al., 2020), scikit-learn 1.6.1 (Buitinck et al., 2013), matplotlib 3.9.3 (Hunter, 2007), seaborn 0.13.2 (Waskom, 2021), and scipy 1.13.1 (Virtanen et al., 2020). As input, a gene expression matrix transformed using the variance-stabilizing transformation (VST) from DESeq2 (Love et al., 2014) was used, with grouping by treatment condition. The clustering of replicates within each group indicates the reproducibility of the RNA-seq experiment**. (B)** Venn diagram showing differentially expressed genes (DEGs) identified at 48 hpi in three comparisons: KO vs WT, KO vs Mock, and WT vs Mock. Genes unique to the KO vs WT comparison (green) were retained for downstream analyses. Venn diagrams were created using Intervene (Khan & Mathelier, 2017). **(C)** Volcano plot of fungal DEGs in maize leaves infected with the Δ*CgEP4* strain compared to WT. Genes significantly upregulated (blue) and downregulated (yellow) are shown (|log₂FC| > 2, adjusted *p* < 0.05), as visualized using the EnhancedVolcano package (Blighe et al., 2018). All plots were generated in R 4.0.2 on macOS.

**Figure S11.**
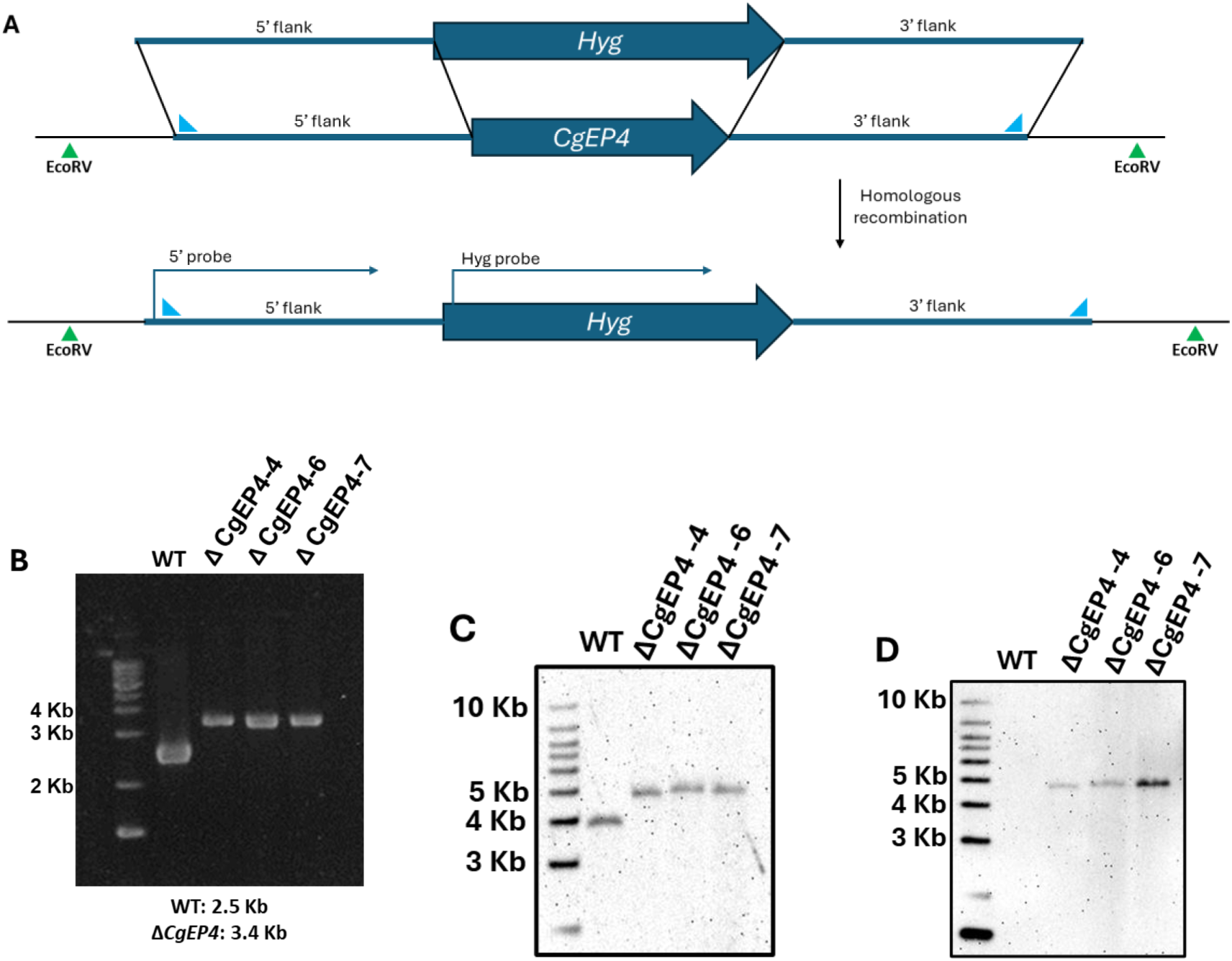
Strategy for deletion of *CgEP4* and molecular validation of transformants. **(A)** Schematic representation of the gene replacement strategy used to delete the *CgEP4* gene by homologous recombination. The *hygromycin resistance* gene (*Hyg*) was flanked by ∼1 kb upstream and downstream sequences of *CgEP4* (5′ and 3′ flanks). Homologous recombination replaces the endogenous *CgEP4* locus with the *Hyg* cassette. Green triangles indicate EcoRV restriction sites used for Southern blot analysis. Blue triangles indicate 9337NesF and 9337NesR primers used to confirm correct integration of the cassette. **(B)** PCR validation of ΔCgEP4 transformants. Primers located within the flanking regions and the *hygromycin* resistance cassette were used to confirm the presence of the deletion construct. A representative gel showing successful amplification in three selected transformants is presented. **(C)** Southern blot using the 5′ flank-specific probe. Genomic DNA from wild-type and mutant strains was digested with EcoRV. The probe hybridized to a 3.7 kb fragment in the wild-type and to a 4.6 kb fragment in the ΔCgEP4 mutant, confirming homologous recombination. **(D)** Southern blot using a Hyg-specific probe. This probe detected no signal in the wild-type, and a single 4.6 kb band in the mutant, indicating single-copy integration of the *Hyg* cassette.

**Figure S12.**
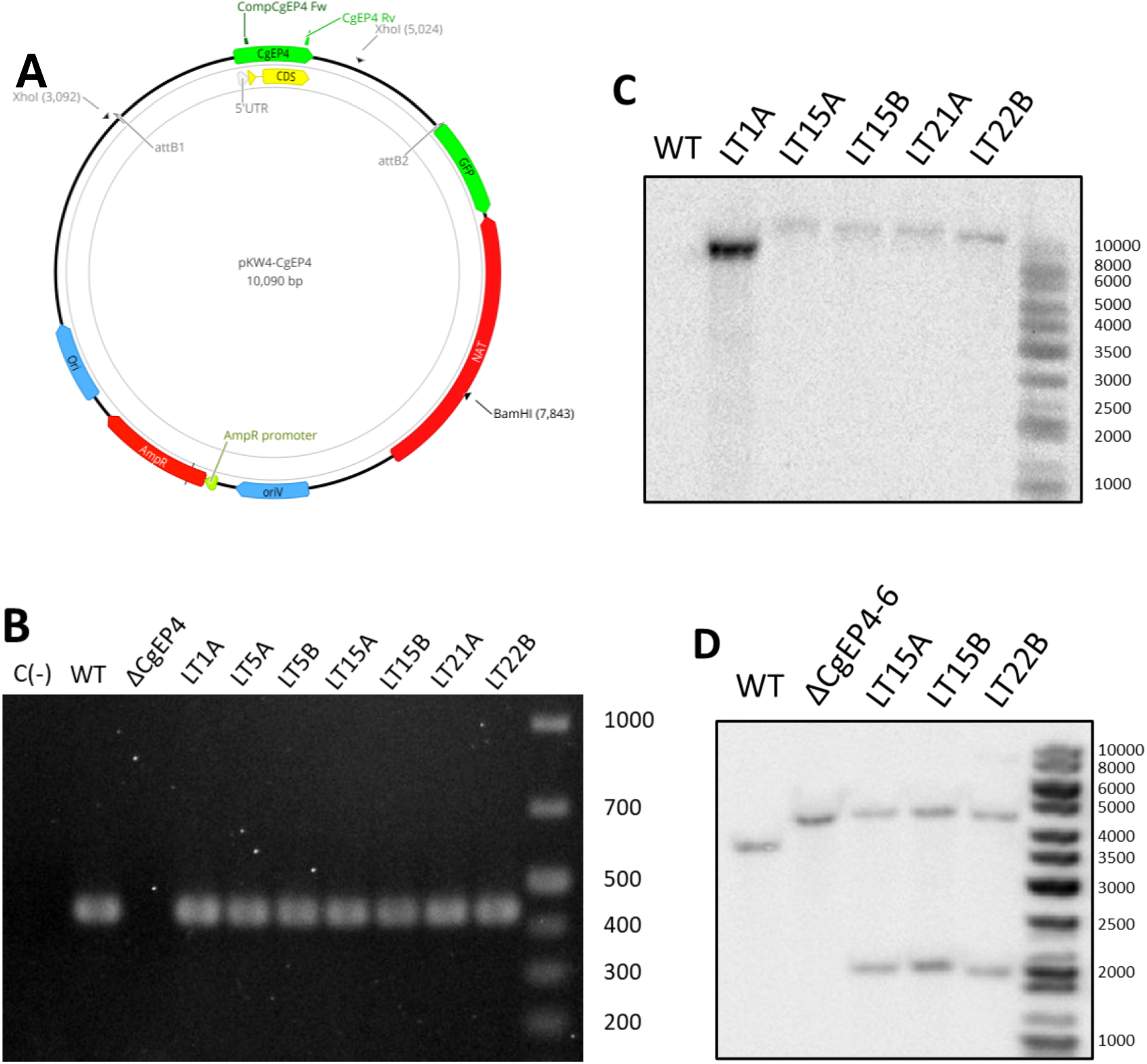
Construction and validation of Δ*CgEP4*::*CgEP4* complemented strain. **(A)** Schematic representation of the complementation construct used to reintroduce the *CgEP4* gene into the Δ*CgEP4* mutant. The coding sequence of *CgEP4*, flanked by approximately 1 kb upstream and downstream regions, was cloned into the pKW4 vector (Vargas et al., 2016), which carries a nourseothricin resistance cassette for selection. Figure generated in Geneious Prime 2025.0.3 (Biomatters Ltd., 2025). **(B)** PCR validation of transformants using primers CompCgEP4Fw and CgEP4Rv to confirm the presence of the *CgEP4* gene. A representative gel showing successful amplification in seven selected transformants is shown. **(C)** Southern blot using a nourseothricin-specific probe. This probe detected no signal in wild-type and Δ*CgEP4* strains, and a single band in complemented transformants, indicating single-copy integration of the complementation cassette. Transformants LT15A, LT15 and LT22B were selected for further validation as they showed a single hybridization band. (D) Southern blot using a 5′ flank-specific probe. Genomic DNA from wild type, Δ*CgEP4*, and complemented strains was digested with XhoI. The probe hybridized to a 3.7 kb fragment in the wild type, a 4.6 kb fragment in the Δ*CgEP4* mutant, and to 4.6 kb and 2.0 kb fragments in the complemented strains. These results confirm that LT15A, LT15, and LT22B are correctly complemented.

**Table S1. CgEP4 and CgEP1 homologs used in phylogenetic analysis.**

**Table S2. *In silico* predictions of CgEP4 subcellular localization.**

**Table S3. Differentially expressed genes in *Zea mays* (Δ*CgEP4* vs WT).**

**Table S4. Differentially expressed genes in *C. graminicola* (Δ*CgEP4* vs WT).**

**Table S5. Primers used in this study.**

**Table S6. *Colletotrichum* species proteomes used in phylogenetic analysis.**

